# Evidence for Transcription and Horizontal Gene Transfer in Dipteran Germline-Restricted Chromosomes

**DOI:** 10.64898/2025.12.11.693641

**Authors:** Riccardo G. Kyriacou, Marion Herbette, Robert B. Baird, Katy M. Monteith, Laura Ross, Christina N. Hodson

## Abstract

Germline-restricted chromosomes (GRCs) constitute a unique class of chromosomes confined to reproductive cells. Arising across multiple evolutionarily distant lineages, GRCs have been identified in three insect families (non-biting midges, gall midges, and fungus gnats), each within the order Diptera. Genomic characterisation in fungus gnats has revealed GRCs to be large, gene-rich chromosomes, which, together with their persistence over millennia, implies they play important biological roles within this clade. However, transcription from these chromosomes has yet to be demonstrated, leaving key questions about their function and activity unresolved. Here, we provide the first direct evidence of GRC-linked gene expression in the fungus gnat *Bradysia coprophila*, integrating RNA-seq data with cytological observations across multiple developmental stages. We report that GRCs express functional genes, though overall transcription is highly limited, likely due to biological factors, such as transcriptional silencing in specific germline cell types, and technical constraints, including filtering to avoid mismapping from core chromosome paralogues. We identify 15 confidently expressed GRC-linked genes using stringent criteria, including five insect homologues of unknown function and nine resembling transposable elements, and report horizontal acquisition of a ∼290 kb bacterial-derived region on GRC2. Furthermore, we perform *in vitro* immunofluorescence staining and confocal microscopy, which indicate increased GRC activity in female oocytes. Overall, these findings establish GRCs in fungus gnats as transcriptionally active, albeit tightly regulated and highly dynamic, chromosomes capable of expressing both endogenous and potentially horizontally acquired genes.

## Introduction

The genetic makeup of multicellular organisms generally remains stable throughout development, with all cells being genetically identical bar mutations. However, it has become increasingly clear that this is not always the case. Programmed DNA elimination (PDE), whereby portions of the genome are systematically removed from specific cell lineages, has evolved repeatedly in diverse metazoans, and even in some single-celled organisms such as ciliates (Bracht et al. 2013; Wang and Davis 2014; Smith 2018). In many of these species, PDE takes place in somatic-fated cells during embryogenesis resulting in germline restricted DNA. The eliminated portion of the genome can range from small DNA fragments to, in some lineages, entire chromosomes known as germline-restricted chromosomes (GRCs). (Wang and Davis 2014; Smith 2018). GRCs have been identified across a large number of diverse, speciose taxa across the tree of life, including hagfish, lampreys, all songbirds and three Diptera families (Kahle 1908; Metz et al. 1926; Bauer and Beermann 1952; Nakai et al. 1991; Pigozzi and Solari 1998). GRCs are distinct from selfish supernumerary (B) chromosomes, nonessential elements that are polymorphic within populations and maintained primarily through meiotic or mitotic drive (Johnson Pokorná and Reifová 2021). In contrast, GRCs appear fixed within populations and have been maintained across deep evolutionary timescales (e.g. ∼50 MYA in songbirds and ∼170 MYA in gall midges) (Lim et al. 2024; Stiller et al. 2024). Their ubiquity and long-term conservation suggest that GRCs might play an important functional role in germline biology (Timoshevskiy et al. 2017; Torgasheva et al. 2019; Hodson and Ross 2021).

In the most well-studied GRC system, songbirds, each species carries a single GRC (Torgasheva et al. 2019; Borodin et al. 2022). Thought to have originated from a selfish B chromosome, given sex biased transmission patterns, the present-day songbird GRC contains multiple copies of genes derived from the core chromosomes (autosomes or sex-chromosomes) (Kinsella et al. 2019; Schlebusch et al. 2023). Many of these genes are actively expressed and enriched in roles in cell division and germline development, perhaps explaining how this selfish chromosome gained important function and thus became domesticated in the germline (Kinsella et al. 2019; Borodin et al. 2022; Vontzou et al. 2023). Comparative data indicates a complex and dynamic evolution of songbird GRC gene retention, with extremely high turnover such that no single gene is shared among all species, despite a common GRC origin (Schlebusch et al. 2023; Ruiz-Ruano et al. 2025). While some genes trace to ancient duplications onto the GRC in the common ancestor of songbirds that were secondarily lost in multiple lineages, others reflect more recent, lineage-specific duplications from the core chromosomes (Schlebusch et al. 2023; Ruiz-Ruano et al. 2025). Notably, only one clear case has been reported in which loss of the core chromosome copy left a GRC-specific gene as the sole homolog (Ruiz-Ruano et al. 2025), consistent with the idea that duplications of genes onto the GRC can allow it to capture essential functions. Subsequent loss of this gene, *zglp1*, from the core genome could render the GRC indispensable and favour stable transmission (Kinsella et al. 2019; Mueller et al. 2023). Once stabilised, aggregation and diversification of genes on the GRC may promote specialisation for germline-specific roles, while avoiding deleterious somatic effects arising from antagonistic pleiotropy (Chapman et al. 2003).

In lampreys, GRC-linked genes are likewise expressed and implicated in gonadal differentiation, oocyte maturation and maintenance, and spermatogenesis (Smith et al. 2012; Yasmin et al. 2022; Timoshevskiy et al. 2025). Analyses of the lamprey germline genome show extensive duplication of core chromosome homologs on GRCs and signatures of positive selection on genes expressed in the germline (Timoshevskaya et al. 2023). These findings indicate that lamprey GRCs, like songbird GRCs, undergo ongoing duplication, turnover, and selection shaping their gene content. Emerging evidence from both clades supports the view that over time GRCs encode important functions, helping explain their long-term maintenance despite rapid gene turnover and the relative paucity of unique, GRC-specific genes (Timoshevskiy et al. 2019; Vontzou et al. 2023; Ruiz-Ruano et al. 2025). Nonetheless, we still lack a comprehensive picture of the evolutionary forces and functional consequences that govern GRC origin, maintenance, and diversification

A striking gap in our understanding of GRC function and evolution exists because most work so far has focused on vertebrate examples. Indeed, the repeated evolution of GRCs in Diptera, the only known insect order to exhibit these chromosomes, remains poorly understood. GRCs are found in at least three highly speciose (Hebert et al. 2016; Kranzfelder and Leonard C Ferrington 2018; Kjærandsen 2022) dipteran families: Sciaridae (fungus gnats), Cecidomyiidae (gall midges) and Chironomidae (non-biting midges) (Hodson and Ross 2021). Dipteran GRCs appear to have originated once in each clade and, like in vertebrates, show evolutionary preservation across each clade, dating back as far as ∼170MYA for gall midges, ∼44MYA for the fungus gnats and ∼120MYA for the non-biting midges (Shin et al. 2013; Hodson and Ross 2021; Lim et al. 2024; Xu et al. 2025). Cytological studies suggest that Cecidomyiidae and most Chironomidae carry many GRCs (up to 80), seemingly as significantly rearranged and highly polyploid copies of the autosomes (Stuart and Hatchett 1988; Staiber and Schiffkowski 2000; Hodson and Ross 2021). In contrast, Sciaridae GRCs are far less numerous and, while containing many paralogous genes to the core genome, appear significantly diverged from the core chromosome set and (for species with multiple GRCs), from each other (Hodson et al. 2025). A key difference with the vertebrate GRCs is that dipteran GRCs comprise a much larger portion of the total genome. In the fungus gnat *Bradysia coprophila*, GRCs comprise almost 40% of the total genome (Hodson et al. 2022). Currently, chromosome level GRC assemblies exist for three fungus gnats (*Bradysia coprophila*, *B. impatiens* and *Lycoriella ingenua*) (Hodson et al. 2025). Comparative analysis across these three species has revealed that the GRCs are generally large, gene-rich (carrying between 3,000-12,000 genes), and for species which carry more than two in their germline (*B. coprophila and L. ingenua*), non-homologus. Another unique distinction of fungus gnat GRCs to those in other taxa is the presence of genes unique to the GRCs, in addition to the many paralogous GRC-linked genes that presumably duplicated onto the GRCs from the core chromosomes (Hodson et al. 2022; Hodson et al. 2025).

The persistence of these chromosomes throughout millions of years, coupled with high gene density and relatively low repeat content, suggest that fungus gnat GRCs are likely functional elements (Hodson et al. 2022; Hodson et al. 2025). However, despite fungus gnat GRCs having a shared origin–likely following an ancient hybridisation event with a Cecidomyiid ancestor–few genes are conserved among those sequenced thus far (Hodson et al., 2025). In addition, studies show that GRCs are often heterochromatic (Metz 1938; Rieffel and Crouse 1966) and display features of selfish genetic elements such as sex-biased transmission and copy number variation throughout development (Gerbi 1986; Gerbi 2022), questioning their functional necessity. Overall, while certain genomic features hint at a potential biological role, parallel to patterns seen in vertebrate GRCs (Schlebusch et al. 2023; Timoshevskiy et al. 2025), the absence of direct evidence for gene expression in dipteran GRCs means it remains uncertain whether these chromosomes serve essential germline functions or persist as largely inert and selfish genomic elements.

There are currently no gene expression studies for dipteran GRCs. Without this data it is difficult to understand the function of these chromosomes; which GRC encoded genes are actually expressed during development, and to interpret the patterns of molecular evolution observed across them. In this study, we analyse gene expression data from germline tissue across four life stages and in both sexes of *B. coprophila* to explore two fundamental questions in regards to function of Scarid GRCs. Firstly, can we detect expression from these two large, gene-rich chromosomes and, if so, what can the types of genes expressed tell us about the function and evolution of GRCs in these flies? Further, we conduct cytological analyses to determine whether GRCs show evidence of transcriptional activity during two stages of development. We find that despite their large size and high gene density, evidence for expression is limited to only a handful of GRC-linked genes, many of which appeared to be transposable elements or insect genes of unknown biological function. We also find horizontal acquisition of a ∼290Kb *Rickettsia*-derived region on one GRC, containing a gene that seems to be transcriptionally active during larval development. We discuss the potential biological and technical reasons for limited expression of GRC genes, and present cytological evidence in testes and ovaries indicating that GRCs, which remain as highly condensed heterochromatic bodies throughout male development, become decondensed and transcriptionally active in female oocytes. These findings suggest that, rather than being broadly transcriptionally active, the GRC may represent a largely heterochromatic and transcriptionally quiescent compartment of the germline genome, punctuated by limited but potentially significant gene activity, which may even include horizontally acquired sequences.

## Results

### GRC-Linked Genes Exhibit Limited but Detectable Expression

To investigate the transcriptional activity of GRC-linked genes, we performed bulk RNA-seq on four life stages in *B. coprophila*: 0-4 h embryos (before GRC elimination), 4-8 h embryos (during GRC elimination), late larval/early pupae, and adults (**Figure 1**). For stages after GRC-elimination from somatic cells (late larva/pupa and adult), both germline and somatic libraries were generated (**Figure 1B**).

**Figure 1.**
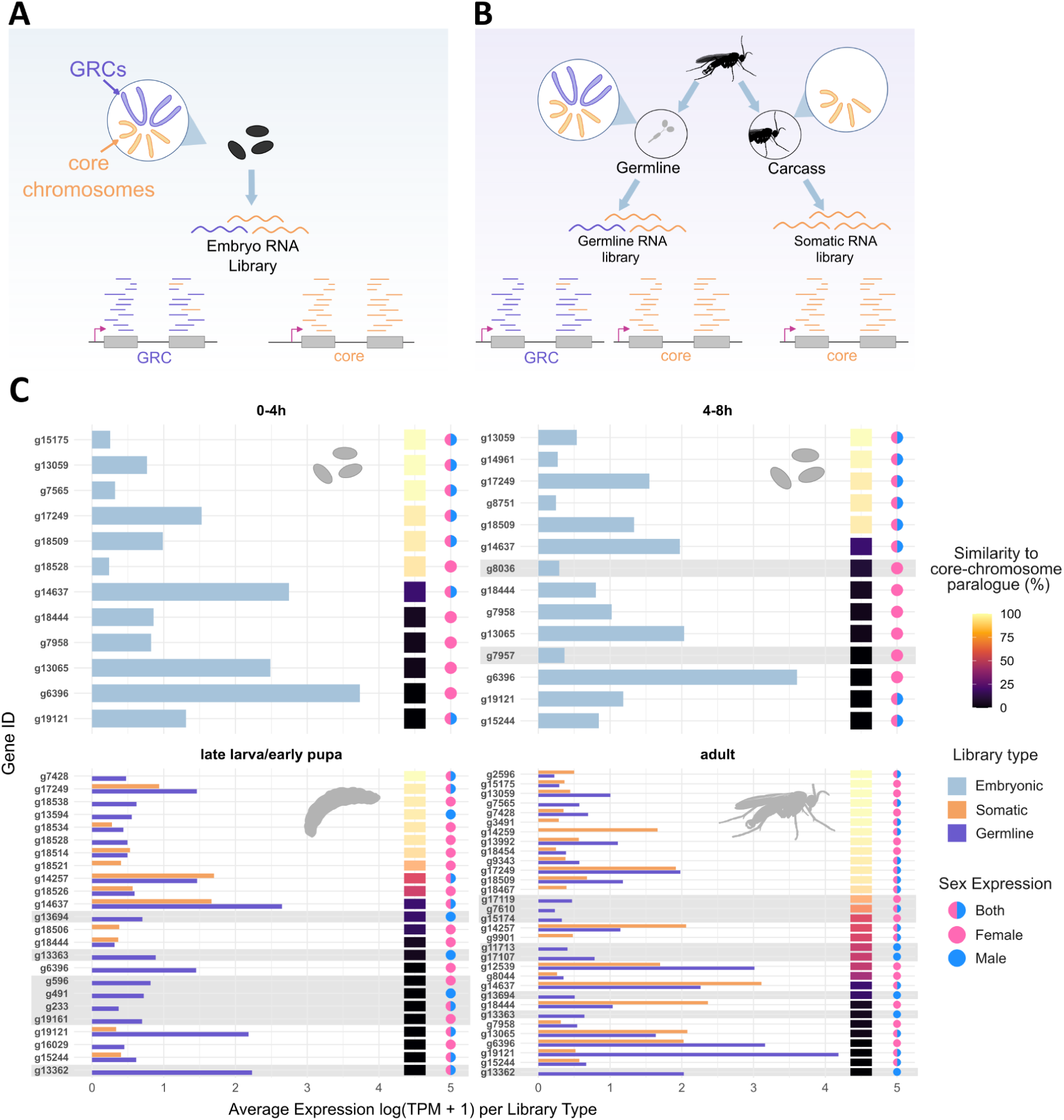
Filtering strategy to identify confidently expressed GRC-linked genes. **A.** Schematic showing the preparation and chromosomal content of the two embryonic RNA-seq libraries (0-4h and 4-8h embryos) utilised in this analysis. Embryonic libraries were prepared before (or during) GRC-elimination, hence all cells contained both core chromosomes (orange) and GRCs (blue). **B**. Schematic showing the preparation and chromosomal content of late larval/early pupal and adult RNA-seq libraries. In these stages, GRC elimination from soma allows for the creation of germline libraries from testes/ovaries and somatic libraries from carcasses. Somatic libraries (right) contain only somatic cells, thus only core chromosomes are present. Germline libraries (left) contain a mix of germline cells retaining both core chromosomes and GRCs, as well as the somatic cells with only core chromosomes. **C**. Plot showing all GRC genes with expression above the significance threshold (y-axis) plotted against their average TPM across libraries in which they were expressed (x-axis). For larval and adult stages, both the TPM in the germline (blue) and somatic (orange) samples are plotted. Each gene is also represented by a coloured bar indicating its sequence similarity to the core genome (calculated as %identity × %coverage), and a circle denoting whether this gene was expressed in males (blue), females (pink) or both (blue and pink). Genes with low core similarity and no detectable expression in somatic libraries at any other life stage were considered confidently expressed (highlighted in grey).

We found 52 GRC-linked genes that exceeded the active TPM threshold (∼0.22) in at least one stage (see methods). However, most of these genes (37) also showed spurious expression in somatic libraries (**Figure 1C**). Notably, this mismapping occurred despite attempts to optimise mapping parameters to reduce multi-mapping and ensure competitive alignment between core (autosomes and X chromosome) and GRC loci. Because GRCs are absent from somatic cells, any apparent expression in these libraries reflects either: (1) misaligned core-derived reads mapping to similar paralogues on the GRCs or (2) germline contamination in the somatic libraries (***Supplementary Figure S2***). In the case of the former, these same misaligned core chromosome reads could falsely inflate expression estimates in germline libraries, making such GRC genes unreliable for this analysis. Hence, genes with observed somatic mismapping and/or significant similarity to core genome paralogues were not considered confidently GRC-expressed. After excluding genes with somatic mismapping or high similarity to core paralogues, only 15 genes remained as confidently expressed (highlighted in grey, **Figure 1C**). This stringent filtering likely underestimates the total number of transcribed GRC genes but yields a high-confidence set given the technical constraints of bulk RNA-seq.

For comparison, 15,296 of 17,181 core chromosome genes were expressed above the same threshold in at least one stage, with broad expression across development and a slight increase in later stages (**Figure 2A**). In contrast, GRC expression is highly limited despite their abundance of coding genes relative to the core genome. Of the 19,917 annotated GRC-linked genes only 52 were expressed prior to filtering, with a total of 85 expression events across all stages (**Figure 2B**). Following our aforementioned filtering strategy to reduce false positives, these numbers drop even further with only 15 confident GRC-linked genes expressed, nine on GRC1 and six on GRC2. Only three of these genes were expressed in more than one developmental stage (**Figure 2C**). These results indicate that, while GRCs are transcriptionally active, their contribution to the transcriptome may be small relative to core chromosomes, particularly when potential false positives are excluded.

**Figure 2.**
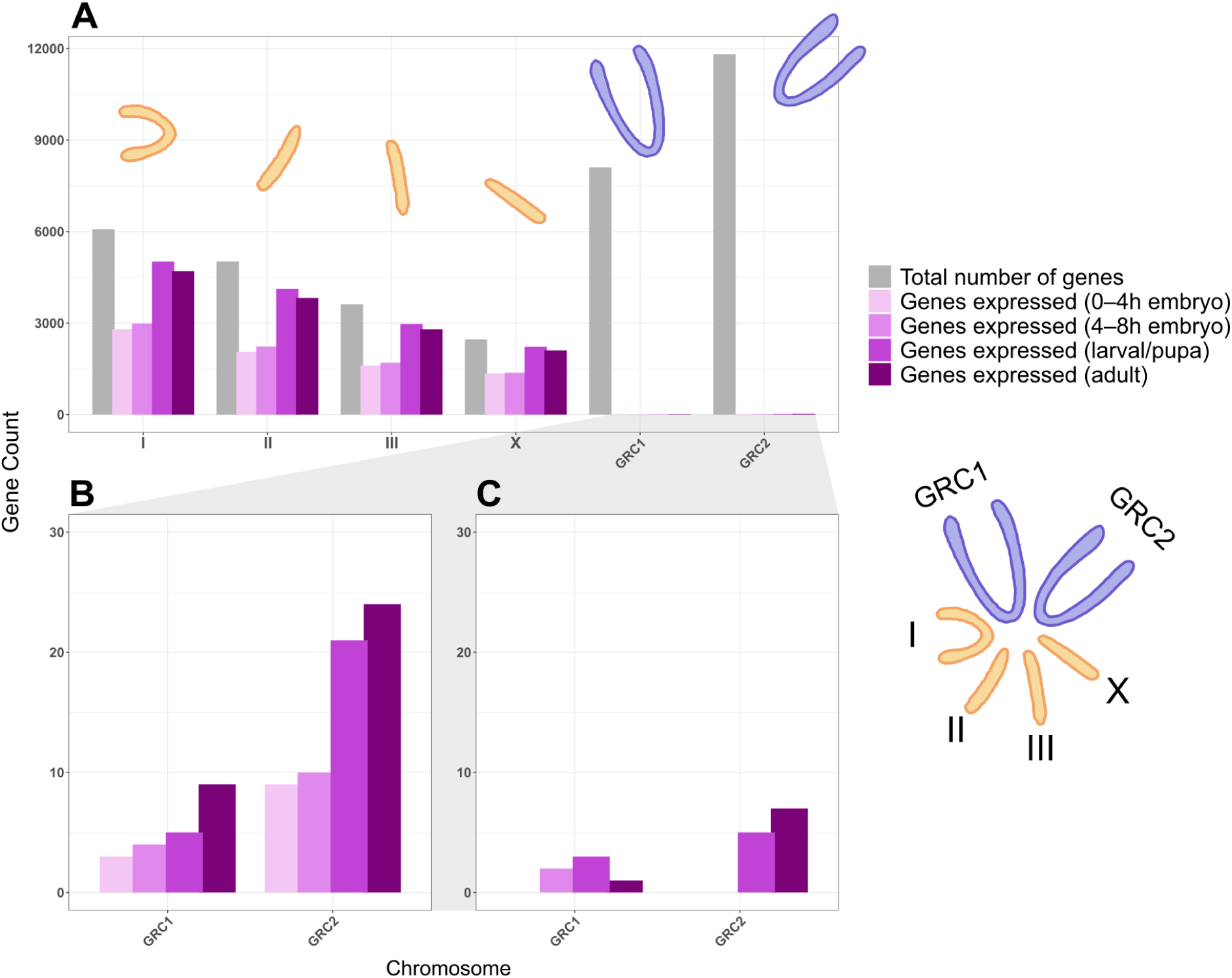
Relatively low but detectable gene expression in *B. coprophila* GRCs. **A.** Plot of gene count (y-axis) against all chromosome scaffolds (x-axis) for B. coprophila. Grey leftmost bars represent the total number of genes annotated for the chromosome. Purple bars represent the number of considered genes expressed per chromosome scaffold, ordered and shaded (lightest to darkest) according to development stage (0-4 h embryos, 4-8h embryos, late larval/early pupa and adult). **B**. Zoomed in plot of only expressed gene counts (y-axis) against GRC scaffolds, again ordered and shaded by developmental stage. **C**. Plot of expressed gene counts (y-axis) against GRC scaffolds, with final counts corrected for mismapping.

Zygotic genome activation (ZGA), the developmental time point whereby the embryo’s own genome initiates transcription, shifting control away from maternally deposited factors, likely occurs when the embryo cellularizes around 12 hours post fertilisation (de Saint Phalle and Sullivan 1996). Since the embryo samples we analysed were <8h post fertilisation, they likely contain almost exclusively maternally deposited transcripts. However, GRC genes could be transcriptionally upregulated after ZGA. In order to account for this potential gap in our data, we additionally analysed RNA-seq data from pooled embryos that were collected between 2h and 2 days post fertilisation (Urban et al., 2021), thus capturing ZGA. We found evidence of 8 GRC-linked genes expressed throughout this broader period of embryonic development, all of which had appeared in our previous analysis (***Supplementary Table S2***). Furthermore, if we remove any expressed GRC-linked genes from Urban et al. (2021) that had shown evidence of mismapping to somatic libraries in our data, only a single GRC-linked gene is found to be confidently expressed. Consequently analysis of bulk RNA-seq data from post-zygotic genome activation embryos does not provide evidence for a limited burst of GRC activity occurring around this timepoint. However, we note that whilst a similar low number of genes were found to be expressed, the TPM values were generally higher for the expressed genes during this time range (5.81 vs 0.57 for the single confidently expressed gene, ***Supplementary Table S2***).

### Sex-Limited Expression of GRC Genes but Comparable Expression Levels Between Sexes

For the GRC genes considered confidently expressed, we examined patterns of expression between sexes. In the 4-8 h embryonic stage, both expressed genes were female-specific (i.e. only expressed in female libraries). In the late larval/early pupal stage, eight genes were expressed: three were female-specific, three were male-specific, and two were expressed in both sexes. In adults, two genes were female-specific, five were male-specific, and one was expressed in both sexes. Overall, these results suggest no strong bias in sex-specific GRC-gene expression by developmental stage, aside from a potential female bias in the earliest embryonic stage (although an extremely small sample size, n=2, must be noted). To formally test differences in expression levels, TPM values were compared between sexes at stages with expression in both males and females using Welch’s t-test. In the late-larval/early-pupal stage, TPMs did not differ significantly between sexes (p = 0.152), whereas in adults, males exhibited significantly higher expression of GRC-expressed genes compared to females (p = 0.0086; ***Supplementary Figure S4***).

These results indicate that GRC-linked genes are expressed in both sexes, although adult males show higher overall GRC-gene expression than females. However, several technical considerations limit interpretation of these comparisons. TPM values, while normalised between libraries, may not accurately reflect true expression levels since germline tissue content likely differs between male and female libraries. Germline libraries contain a mixture of somatic and germline cells, and male libraries may have a higher germline-to-soma ratio than female libraries, potentially biasing apparent expression levels. Given these technical limitations and the limited statistical support across developmental stages, we conclude that our RNA-seq data provide no robust evidence for sex-biased GRC transcription. Nonetheless, most GRC-linked genes (12 of 15) appear to be expressed exclusively in one sex.

### Expressed GRC-Linked Genes Exhibit Homology to Selfish Genetic Elements and Uncharacterised Insect Proteins

To investigate the potential functions of the 15 GRC-linked genes confidently expressed across *B coprophila* development, homology-based analyses at the protein level were performed (**Figure 3A**). Gene names in this study correspond to their names in Hodson et al. (2025) (see methods).

**Figure 3.**
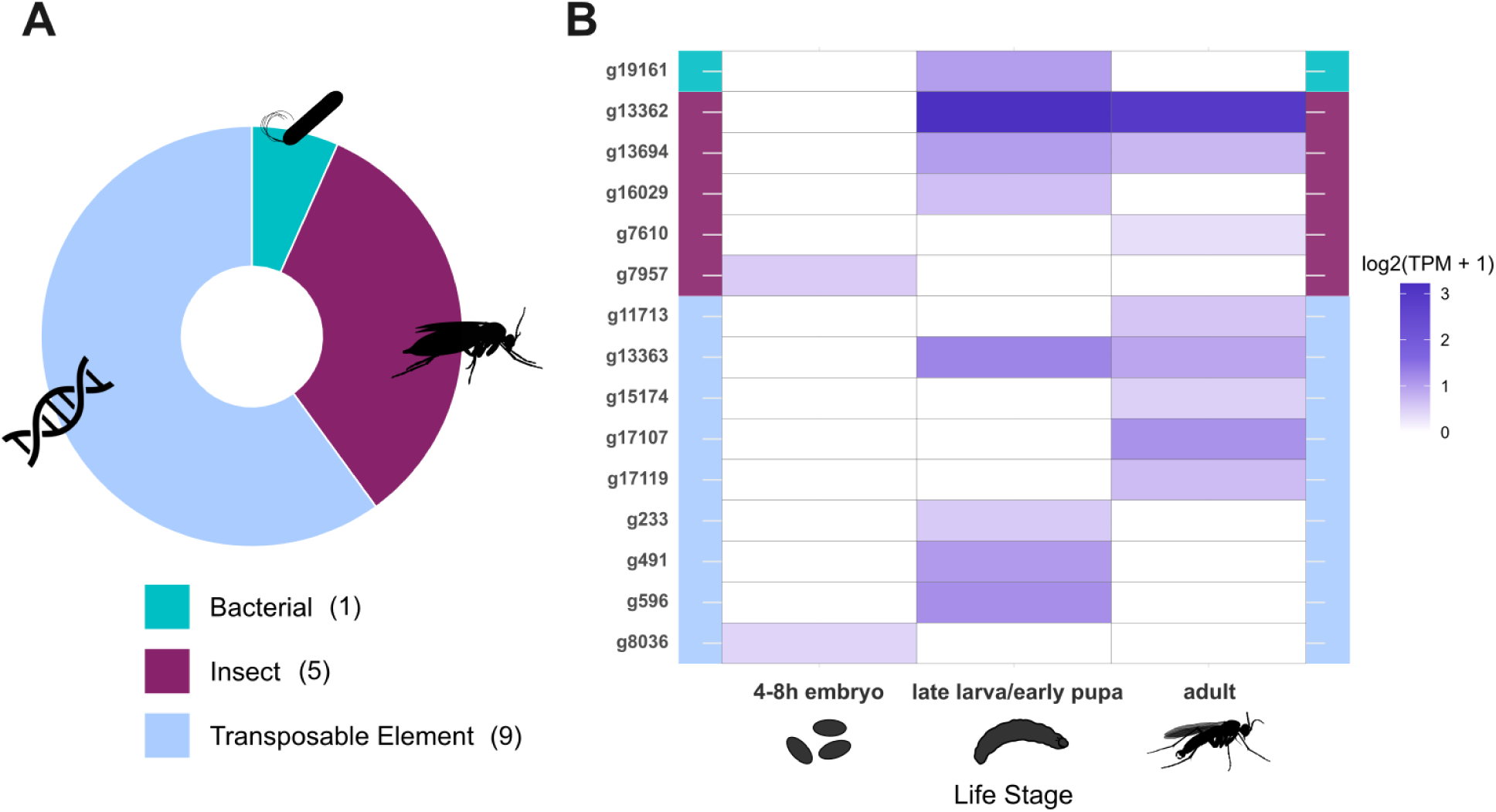
Inferred origin and relative expression levels for 15 GRC-linked genes. **A.** Pie chart showing the number of GRC-linked genes (n = 15) assigned to each top BLAST hit category. Each gene was classified based on its highest-scoring BLAST hit (against the NCBI non-redundant protein database and Repbase), and categorised as a transposable element, insect, bacterial, or no detectable homology. Nine genes were classified as transposable elements (g233, g491, g596, g8036, g11713, g13363, g15174, g17107, g17119), five as insect genes (g7610, g7957, g13362, g13694, g16029) and one as bacterial (g19161). **B.** Heatmap showing expression levels (log₂(TPM + 1)) of the same GRC-linked genes across developmental stages. Genes are ordered by homology category (from top to bottom: Bacterial, Insect, Transposable Element), with a side colour bar indicating category. Expression values were averaged across all available sex and tissue libraries for each gene and developmental stage.

Nine of the expressed GRC-linked genes showed homology to transposable elements (TEs). Of these, five were similar to retrotransposon families: *BEL* (*g11713, g17107*), *CR1* (*g15174*), *Copia* (*g17119*), and Vingi (*g8036*). Four additional genes exhibited homology to DNA transposons: *Kolobok* (*g13363*), *piggyBac* (*g233*, *g491*), and *Transib* (*g596*).

Five GRC-linked genes displayed homology to insect proteins. Four of these *(g13694, g16029, g7610, g7957*) showed homology to “hypothetical proteins” of unknown function identified in other species. Further analysis of inferred conserved protein domains revealed that *g16029* encodes a SET domain–containing protein typically associated with histone methyltransferase activity, while *g13694*, *g7610*, and *g7957* lacked identifiable domains. One gene, *g13362*, showed partial BLASTp homology to *Lithostathine-2* from the fungus gnat *Pseudolycoriella hygida* and was relatively strongly expressed, compared to other GRC-expressed genes, during the later developmental stages (**Figure 3B**).

One GRC-linked gene, *g19161*, located on GRC2, showed strong homology to a bacterial gene. BLASTp identified it as homologous to the nucleotide exchange factor *GrpE*, a conserved chaperone cofactor, in various bacterial species. Notably, this includes *Rickettsia prowazekii*, an endosymbiotic bacterium known to inhabit the germline cells of *B. coprophila* (Gutzeit et al. 1985; Urban et al. 2021). This gene is also present in the *Rickettsia* cobiont assembly generated when decontaminating the *B. coprophila* genome. This finding led us to investigate whether there was a horizontal gene transfer event from the cobiont onto GRC2.

### Long-read Evidence Supports Integration of *Rickettsia*-Derived *GrpE* into GRC2

The homology between *g19161* and *Rickettsia prowazekii GrpE* suggests two possibilities: (1) integration of *GrpE* into GRC2 via horizontal gene transfer (HGT) from a bacterial endosymbiont, or (2) erroneous assembly of a bacterial contig, originating from *Rickettsia*, into the *B. coprophila* genome assembly. To distinguish between these scenarios, the raw PacBio long-read sequencing data, originally used to assemble the *B. coprophila* reference genome, was re-mapped back to the annotated genome.

The alignment revealed that *g19161* resides in a ∼290 kb region on GRC2 with substantially elevated read coverage compared to the rest of the genome (**Figure 4**). Notably, all 101 genes within this region (*g19067-g19167*) show strong homology to bacterial proteins, many from *Rickettsia*, suggesting a likely bacterial origin. The elevated coverage across this segment is consistent with the presence of bacterial DNA in the original genomic DNA libraries used for assembly. A visible dip in local GC content, typical of bacterial DNA, is also visible across the bacterial-region, from the GRC2 average of 36% to 34% (**Figure 4A**). Importantly, this region is flanked by genes of clear insect origin: two highly conserved single copy insect genes are located just outside the high-coverage, lower GC segment: *g19053* (*General transcription factor IIH subunit 1*) lies a few genes upstream, and *g19168* (*ectopic P granules protein 5 homolog*) is located immediately downstream (**Figure 4A**).

**Figure 4.**
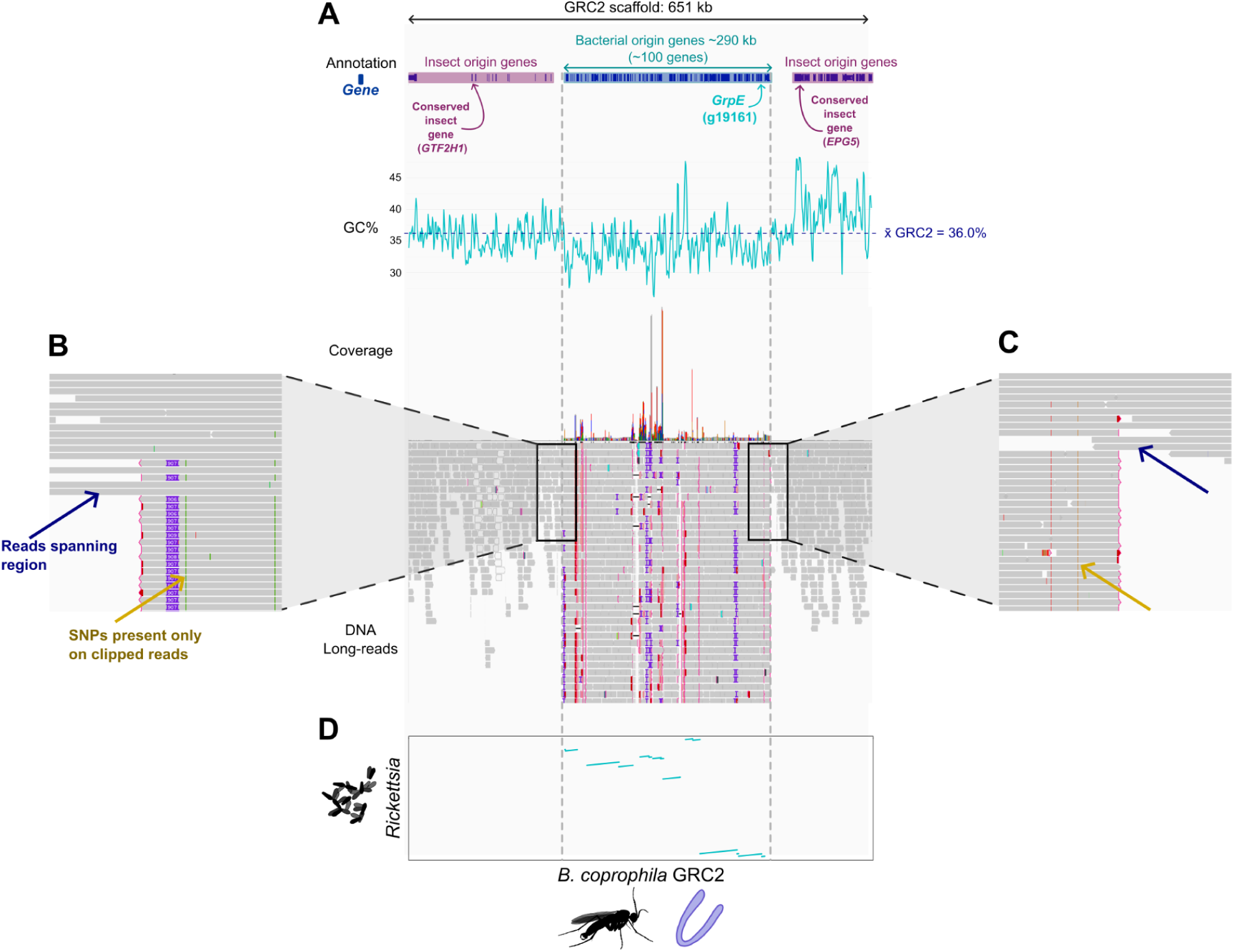
GRC-linked gene *GrpE* exists in a putative HGT region consisting of ∼100 bacterial-origin genes. **A.** 651kb section of GRC2 containing ∼290kb bacterial-derived region. Putative HGT region boundaries highlighted by grey vertical dotted lines. Insect origin genes flanking the HGT region highlighted in pink, including two conserved (BUSCO) genes: *GTF2H1* and *EPG5*. Bacterial origin genes highlighted in blue, arrow pointing to g19161 (GrpE). Below, GC content (%) in a 2kb sliding window is plotted across the 651kb region. Mean GC content for the whole of GRC2 (dark blue dotted line, 36.0%) superimposed, calculated across a 200kb sliding window. Visible drop of GC content seen for putative HGT region (mean GC = 34.1%). Below, integrated genomics viewer (Thorvaldsdóttir et al., 2013) track showing read coverage graph and Pac-Bio long read alignments across the region. **B.** Zoom in showing the leftmost HGT-region and read coverage boundary. Blue arrow (top) points to long reads spanning across the high-coverage boundary (n=15). Yellow arrow (below) points to more diverged reads abruptly clipping at the boundary (soft-clipping >100bp highlighted as red line at the end of read), all containing SNPs not present in flanking reads. **C.** Rightmost zoom into HGT-region and read coverage boundary showing reads spanning across the boundary (n=10). Yellow and blue arrows denote the same as B. **D.** Dotplot showing DNA alignment between the 651kb section of GRC2 and the co-assembled endosymbiont *Rickkestia* genome. Alignments between the genome occur exactly within the putative HGT region denoted via the grey vertical dotted line.

Visual inspection of long-read alignments across the flanking boundaries reveals multiple full-length reads that span the bacterial-homologous region and extend into adjacent, lower-coverage GRC2 sequence (**Figure 4B/C,** blue arrows). These spanning reads (15 downstream and 10 upstream of the putative HGT region; ***Supplementary Figure S5***) support the physical integration of this region into the GRC scaffold. In contrast, most reads mapping to the high-coverage region are truncated at the boundaries, exhibiting soft clipping (>100 bp) and numerous SNPs not present in the spanning reads (**Figure 4B/C**, yellow arrows). These diverged, clipped reads likely originate from bacterial DNA present in the original genomic libraries and, notably, all terminate precisely where the GRC-derived sequence begins. Together, these patterns strongly support the conclusion that this bacterial region is physically integrated into GRC2, rather than representing contamination or misassembly.

In order to test whether this integrated bacterial region has derived from endosymbiotic Rickettsia known to exist within the germline (Urban et al., 2021), we performed a genome alignment between GRC2 and the *Rickettsia* genome co-assembled with the *B. coprophila* assembly. Dot-plot visualisation (**Figure 4D**) revealed strong alignment between the two species occurring across the HGT region and terminating precisely at the boundaries of the extremely high coverage region. Hence, we suggest that the ∼290kb bacterial region present in GRC2 has been integrated though HGT with an endosymbiotic *Rickettsia*. However, whilst sequence similarity between these two regions was well generally conserved (with only ∼150 SNPs occurring between the ∼291 kb of aligned sequences), synteny is shuffled, suggesting substantial genomic rearrangements occurring in this region of GRC2 post-integration.

### Cytological Evidence for GRC Transcription

To complement our RNA-seq analysis, we performed immunofluorescence staining on *B. coprophila* testes and ovaries from late larval (4th instar) and adult stages. This analysis is aimed to assess the transcriptional activity of GRCs by examining chromatin states and histone modification patterns. Embryonic stages (0-4h and 4-8h) were excluded from staining because zygotic genome activation has not yet occurred at these time points. Consequently, while maternally deposited GRC transcripts can be detected computationally, active GRC transcription is absent, rendering such immunofluorescence staining uninformative for these early embryo stages. Tissues were stained with an antibody against H4K16ac, an evolutionarily conserved histone modification associated with euchromatin and active transcription, along with Hoechst for DNA visualisation (**Figure 5**).

**Figure 5.**
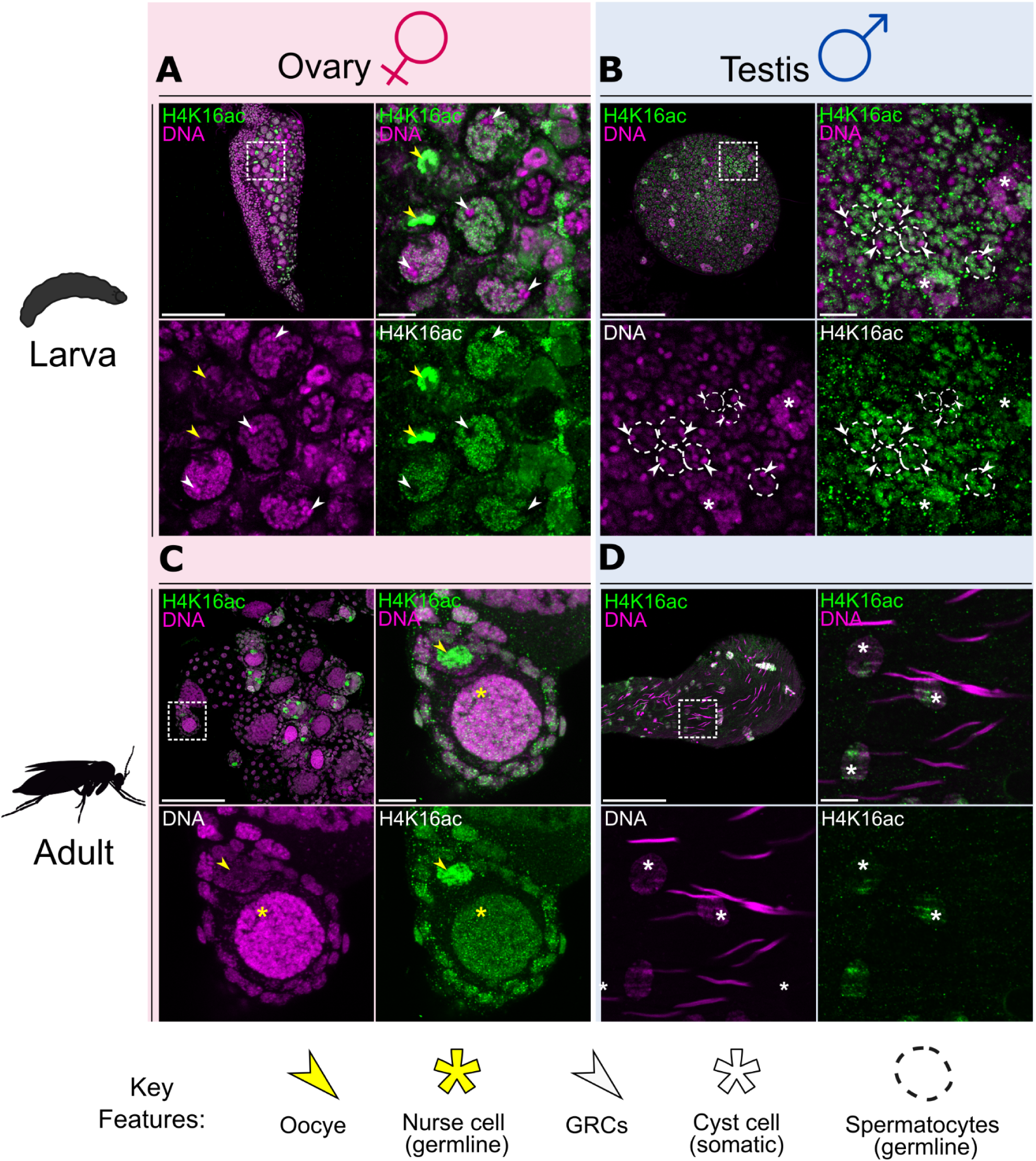
Confocal images of whole-mount testes and ovaries from *B. coprophila* adults and larvae stained with anti-H4K16ac antibody (green) and Hoechst for DNA (purple). **A.** larval ovary, with an overview and magnified region showing oocytes (yellow arrowheads) brightly stained for H4K16ac and polyploid nurse cells with highly condensed and non-acetylated GRC chromosomes (white arrowheads). **B.** larval testis, with an overview and magnified region showing primary spermatocytes and somatic cyst cells (asterisks), a couple of individual primary spermatocytes are outlined (dashed lines), white arrowheads indicate condensed, non-acetylated GRC chromosomes which are only present in primary spermatocytes. **C**. adult ovary, with an overview and magnified region of nurse cells (asterisks) with low levels of H4K16ac and an oocyte (yellow arrowhead) brightly stained for H4K16ac, both lack condensation of the GRCs. **D**. adult testis, with an overview and a magnified, showing needle-shape sperm nuclei and round somatic cyst cells (asterisks), mature sperm cells lack detectable H4K16ac likely because they underwent replacement of histones by protamines. Scale bar: 100µm (overview images) and 10µm (magnified regions).

Immunofluorescence staining and confocal microscopy revealed that in some cell types/ developmental stages, GRCs are highly condensed into heterochromatic bodies, potentially explaining the limited transcription detected via RNA-seq. In male larvae, testes staining sees a repeated clustering of two highly condensed chromosomes (intense magenta foci; **Figure 5B**) which are absent of H4K16ac staining, and appear in contrast to decondensed, highly acetylated chromosomes (likely the core chromosomes). These condensed bodies, present only in the primary spermatocytes, are very likely the GRCs, and previous cytological reports have reported such GRC decondensation in *B coprophila* larvae of a similar developmental stage (Rieffel and Crouse 1966). In adult male testes, we see no H4K16ac in mature sperm (needle-shape nuclei, **Figure 5D)**, which is expected due to the histone-to-protamine transition that renders chromatin completely inaccessible by the polymerase machinery (Rathke et al. 2014).

In female larvae, confocal microscopy revealed two distinct cell populations with differential staining patterns. Small cells exhibited uniform DNA staining and widespread H4K16ac signal, consistent with globally euchromatic and transcriptionally active chromosomes (yellow arrowheads, **Figure 5A**). In contrast, larger cells contained multiple intensely DNA-stained, condensed chromosomes void of H4K16ac, indicating transcriptional repression. Based on size, morphology, and comparison with previous studies in *B. ocellaris*, the small cells likely correspond to primordial germ cells, while the larger cells are polyploid nurse cells (Berry 1941). Both cell types are germline-fated and thus should contain GRCs in females. In primordial germ cells, the presence of uniformly euchromatic chromosomes and strong H4K16ac signal suggests that GRCs are transcriptionally active (Goday and Ruiz 2002; Taylor et al. 2013; Zhang et al. 2021). On the other hand, in nurse cells, we observe highly condensed, transcriptionally inactive chromosomes (white arrowheads, **Figure 5A**) appearing in groups of four, consistent with polyploidy and heterochromatinisation of GRC1 and GRC2. Female adult ovaries saw no condensed chromosome bodies in either the nurse cells nor the oocytes. This along with ubiquitous H4K16ac staining throughout the euchromatic nuclear environment suggest all chromosomes, including GRCs, are decondensed and transcriptionally active.

In summary, immunofluorescence staining and confocal microscopy in male and female germline tissue support a model of limited GRC transcriptional activity, due to differential chromatin/histone states dependent on sex and cell type. In males, GRC transcription appears highly limited. In females, larvae see GRCs being transcriptionally active in developing oocytes (primordial germ cells) but become heterochromatic and silenced in polyploid nurse cells. Adult female staining suggests decondensed and actively transcribing GRCs in both nurse cells and oocytes.

## Discussion

### *B coprophila* GRCs Transcribe Genes, but Expression is Limited

RNA-seq data from this study suggests only a limited set of GRC-linked genes are expressed, despite the relative gene richness of these chromosomes. This finding could have multiple explanations. One biological factor may be that GRCs often appear transcriptionally repressed, with older cytological studies noting that GRCs exist as condensed in heterochromatic bodies during some development stages (Metz 1938; Rieffel and Crouse 1966). Immunofluorescence performed in this study expands on these previous observations, suggesting that GRC activity and chromatin state (inferred through staining for the euchromatic histone modification H4K16ac) varies across germline cell lineage and between sexes (**Figure 5**). In female larvae, GRCs appear transcriptionally active and decondensed in oocyte precursors but repressed in polyploid nurse cells, a clear example of differential GRC activity due to germline cell type. Differential GRC chromatin states between sexes can be seen in male larvae of the same stage, where all GRCs visualised remain condensed and heterochromatic. Likewise, in adult females, GRCs appear decondensed and transcriptionally active in both oocyte and nurse cells, while adult male testes see extremely low H4K16ac staining due to the histone-to-protamine transition. Such results would also suggest a clear female bias in adult and larval RNA-seq data, but we find no evidence for this in the data obtained in this study. Specific sex biases aside, if GRCs throughout development often exist in these opposing states of active and inactive transcription, it is not surprising that expression detected from bulk RNA-seq is low relative to the core chromosomes.

However, despite our cytological staining and RNA-seq data, we argue it remains unclear to what extent limited GRC-expression represents a true biological feature of Sciarid GRCs. This is due to two technical considerations that arise with the methods presented in this study that may result in an underestimate of GRC transcription. Firstly, the use of bulk RNA-seq from whole germline tissues introduces a bias that disproportionately favours transcripts derived from core chromosomes over those from the GRCs. Ovaries and testes, from which we established germline RNA-seq libraries, are composed of a mixture of germline cells (containing both core chromosomes and GRCs) and surrounding somatic cells (which contain only core chromosomes) (Du Bois 1932). If, as germline DNA sequencing suggests, the proportion of somatic to germline cells is approximately 1:1 in germ tissue dissections (Hodson et al., 2025), then the overall chromosomal content across all cells in the germline tissue would reflect a ratio of four core chromosomes for every GRC (in diploid cells, see **Figure 1B**). Consequently, this ratio imbalance may lead to core chromosome transcripts “swamping” out GRC-derived transcripts, skewing detected expression away from GRC-linked genes. Indeed, this is a common problem with bulk RNA-seq datasets which contain a mix of gonadal and somatic tissue, but the presence of GRCs makes this issue easily detectable. A second bias limiting detected GRC RNA-seq expression is more intentional, stemming from conservative post-mapping filtering applied to the expression data. This was done to eliminate false-positive detection of GRC-linked genes that may arise from core chromosome derived reads mapping to paralogous GRC sequence, an issue noted in RNA-seq of vertebrate GRCs (Schlebusch et al. 2023). For the purposes of this study where we aimed to confidently demonstrate true GRC expression, it was necessary to account for mismapping in this analysis. Consequently, only genes which showed no signs of mismapping at any life stage (that is, the mapping of reads to the GRC scaffold in somatic, non GRC containing libraries) and had no similar GRC-derived paralogue, were considered expressed. In this way, 37 of the ∼50 GRC-linked genes expressing TPMs above the expressed threshold were disregarded. Not only does this conservative method filter exacerbate our bias in detecting few GRC genes, but it necessarily will only allow for the detection of GRC genes that were not recently derived (and therefore exist as highly similar paralogues) from the core chromosomes.

Overall, these results should be viewed more as a confirmation of GRC-expression, even when imperfect RNA-seq methods and extremely conservative thresholds are employed, rather than a quantitative report on relative GRC expression levels. Future studies employing single cell RNA-seq or single-nuclei cell sorting, to enrich or isolate germline cells, will be able to provide more fine scale analyses of GRC gene expression. These methods will help alleviate issues, particularly in regards to core chromosome contamination and over-representation, and allow for more quantitative assessments of expression. The use of RNA FISH on GRC-linked genes would also eliminate the need for strict computational filtering, and would provide a definitive assessment of which genes are truly transcribed from the GRC. Finally, cytological staining could also be performed on further life stages, giving a more complete time-series of GRC activity throughout development, and potentially highlight stages with punctuated high GRC expression.

### GRCs Express a High Proportion of Selfish DNA

Expression of five non-TE genes in *B. coprophila* GRCs suggests that these chromosomes do have a biological function, potentially explaining their maintenance across populations. However, given the nearest homologues for four of these genes (*g13694, g16029, g7610, g7957*) are annotated only as “hypothetical proteins”, the precise phenotypic effects of GRC transcription remain enigmatic. The only gene with an informative BLASTp result, *g13362*, was a partial homologue of *Lithostathine-2*. Related to the *regenerating gene I alpha* (*REG1A*) in humans, which has pancreatic and tissue repair functions in humans (Mao et al. 2021), *Lithostathine-2* has no described function in insects. Furthermore, none of these five genes had homologues detected in the GRCs of the two other fungus gnats sequenced (*B. impatiens* and *L. ingenua*), implying the high levels of gene turnover reported in Hodson et al. (2025) applies to putatively functional genes (***Supplementary Table S3***). Biologically relevant genes varying rapidly between the GRCs of closely related species occurs likewise in the songbirds (Schlebusch et al. 2023; Ruiz-Ruano et al. 2025), potentially hinting convergent evolutionary forces driving patterns of loss and gain in these unique chromosomes. Further advancing our understanding of GRC function will require the discovery of additional transcribed GRC-linked genes and improved annotation/domain prediction pipelines, with an eventual goal towards using gene editing approaches to empirically assess the function of candidate loci.

Nine of the 15 confidently expressed GRC-linked genes identified in this study showed strong sequence homology to transposable elements (TEs). Although GRCs in the three fungus gnat species sequenced to date appear to be depleted in TE content relative to the core chromosomes (Hodson et al. 2025), several TE families (LINE, LTR, and DNA transposons) are notably enriched on these chromosomes. All nine TE-related transcripts detected in this study belong to these families. Four of these TEs show homology to elements present on GRCs of the closely related fungus gnats *B. impatiens* and *L. ingenua* (***Supplementary Table S3***). Together, our results raise the question of why these specific TE classes accumulate and are transcriptionally active on the GRC, despite an overall depletion of TE sequences at the genomic level. If the GRCs of *B. coprophila* are largely non-recombining, a feature that remains unresolved (Hodson et al. 2022; Hodson et al. 2025), then reduced efficacy of purifying selection could facilitate the accumulation of TEs. Empirical analyses and theoretical models both indicate that the absence of recombination enhances the effects of Hill–Robertson interference, thereby limiting the efficacy of purifying selection against transposable element insertions (Bartolomé et al. 2002; Dolgin and Charlesworth 2008). However, such effects should generally influence all TE families, not just the subsets enriched on the GRC, suggesting that additional, family-specific factors such as insertion-site preference, differential epigenetic regulation, or autonomous replication capacity may contribute to this pattern (Rizzon et al. 2002; Kent et al. 2017; Bourgeois et al. 2020; Catlin and Josephs 2022).

Alternatively, the elevated representation of TE transcripts in our dataset may primarily reflect their increased transcriptional activity in germline tissue. TEs are often upregulated in the germline, where their expression facilitates vertical transmission to subsequent generations (Haig 2016). Given the limited number of protein-coding genes expressed from the GRC (as suggested by our RNA-seq and cytological data), this germline-specific TE activity could disproportionately contribute to the detected transcript pool. Moreover, bulk RNA-seq approaches are inherently biased toward highly expressed loci, further amplifying the relative abundance of TE-derived transcripts in our dataset. Finally, observational bias may also play a role in the TEs found expressed in this data. Because our stringent filtering strategy removed any annotated GRC-linked genes with similar homologues in the core chromosome, any shared TEs between core and GRC with little divergence will not have been considered. Consequently, the use of single-cell RNA-seq approaches, along with a more complete understanding of the TE landscape and expression profiles in the GRCs and core chromosomes, will allow for a greater contextualisation of the results presented in this study.

### Insights into GRC Evolution from Expressed GRC-linked Genes

All expressed GRC-linked genes identified in this study are either unique to the GRCs, or are highly diverged from their core chromosome copies, a factor due to technical considerations minimising false positive discovery rates. Based on this divergence and the presumed hybrid origin of GRCs from Cecidomyiidae (Hodson et al. 2022; Hodson et al. 2025) we consider three possible evolutionary origins for these loci. (1) These genes are Cecidomyiid in origin, having been retained since the introgression, (2) they represent core chromosome genes that duplicated onto the GRC, with the ancestral core copy being subsequently lost, or (3) these genes represent extremely diverged core chromosome copies. These scenarios have different implications for how sciarid GRCs originated and were maintained over the ∼50 million years since the inferred introgression.

If these expressed genes are in fact cecidomyiid in origin, and thus have remained functional for millions of years, their persistence suggests that they contribute to important biological processes in fungus gnats. In this scenario, the presence of functional cecidomyiid genes on nascent sciarid GRCs may have conferred an immediate selective advantage to GRC-containing ancestors, promoting their initial spread and maintenance. Alternatively, the early evolution of GRCs may have been largely neutral, with emergence tolerated by the absence of strong deleterious effects. Subsequent duplication of core chromosome genes with essential functions onto the nascent GRCs could then have provided a route to obligate GRC inheritance and maintenance over evolutionary time. This could have occurred through loss of the ancestral core chromosome copy (as proposed for songbird GRCs; Ruiz-Ruano et al. 2025), or through relaxed functional constraint enabling rapid neo-/subfunctionalisation of the GRC paralogue, ultimately allowing it to acquire essential *de novo* function (Sandve et al. 2018). Regardless of the specific origin, the discovery of highly diverged, GRC-specific genes that are actively transcribed in *B. coprophila* provides new insight into the early evolution of these chromosomes and motivates further functional and comparative genomic work.

### Integration of ∼100 Rickettsia Genes into GRC2 via Horizontal Gene Transfer

Our study also reveals a potential third origin for GRC-linked genes beyond introgression from cecidomyiids and duplication from the core genome: horizontal gene transfer (HGT) from non-insect donors. We identify a ∼290 kb prokaryotic-derived region on GRC2 that contains 101 bacterial genes. This region is unlikely to represent a misassembly artifact, as multiple long reads (which represent a single DNA molecule) span both the bacterial-derived sequence and adjacent insect-like regions, confirming their physical linkage within the chromosome (Suzuki et al. 2019; Bellas et al. 2023). The region shows highly contiguous alignment to the coassembled *Rickettsia* genome, a germline-associated endosymbiont of *B. coprophila* (Urban et al., 2021), indicating *Rickettsia* as the probable donor lineage. Within this putative HGT block, only a single gene, *g19161*, meets our criteria for confident GRC-linked expression (see Results). The GRC2 and bacterial copies show 100% sequence identity, preventing us from confidently determining whether transcripts originate from the GRC or from the endosymbiont. However, it is notable that *g19161* expression is restricted to late-larval samples and occurs at TPM values comparable to other GRC-linked genes. In contrast, bacterial transcripts typically appear across multiple developmental stages and at far higher levels, consistent with the high abundance of bacterial genomes in germline tissues (Gutzeit et al. 1985; Urban et al. 2021). Indeed, other genes within the same bacterial-derived region show stage-wide expression, with one clearly bacterial gene expressed at ∼40 TPM, compared to ∼0.5 TPM for typical GRC-linked genes (***Supplementary Figure S6***). The origin of *g19161* expression therefore remains uncertain. BLASTp analysis identifies strong homology to *Rickettsia* GrpE, a conserved bacterial chaperone cofactor (Harrison 2003) with no known function in insects, further complicating its interpretation.

Nonetheless, the presence and possible expression of bacterial genes within GRC2 suggest that *B. coprophila* GRCs may not only accumulate endogenous duplicates from core chromosomes but also retain horizontally acquired genes. HGT has previously been identified in *B coprophila*, with genes originating from bacteria (Chen et al. 2024; Hoile et al. 2025) as well as springtails and fungi (Feyereisen et al. 2023; Kapoor et al. 2025) having been reported. The latter example of HGT also seems to have occurred in the GRCs themselves, with Feyereisen et al. (2023) reporting the presence of eight GRC-linked genes of “alien” origin. The authors go on to speculate that the extensive chromosome rearrangements and gene translocations seen in fungus gnat GRCs (Hodson et al. 2022; Hodson et al. 2025) may predispose GRCs to HGT from organisms sharing similar ecological niches. Our reported bacterial HGT region supports this idea twofold. Firstly, *Rickettsia* exists as an endosymbiont within the germline cells of *B. coprophila*, and secondly, genomic rearrangements even within this HGT region itself are observed. Overall, our results add to a growing body of literature suggesting HGT may be implicated in the evolution of GRCs, and highlights the GRCs potential role as dynamic platforms for genomic innovation, containing genes from taxonomically related species (e.g. cecidomyiids) via hybridisation, the core genome of sciarids, as well as from distant taxa through HGT. Broader analyses of HGT prevalence across GRCs from multiple fungus gnat species will help clarify whether this ∼290 kb insertion represents a rare event or part of a wider pattern of endosymbiont-derived gene acquisition, providing insights into how horizontal transfer shapes sciarid GRC evolution.

### Conclusion

In conclusion, RNA-seq and immunofluorescence staining across developmental stages reveal that the GRCs of *B. coprophila* exhibit detectable gene expression. To our knowledge, this represents the first direct evidence of active GRC transcription in Diptera. Our observations also contribute to the growing body of work challenging early hypotheses that Sciaridae GRCs are purely selfish chromosomes escaping paternal genome elimination–the unique sex-determining mechanism in fungus gnats by which males transmit only maternally derived chromosomes (Haig 1993; Goday and Esteban 2001). Interestingly, despite GRCs containing more genes than the combined core chromosome set, we find transcription to be highly limited, with only 15 expressed genes being unambiguously GRC-linked. Cytological imagining also reveals differential GRC regulation between sex and germ cell lineages, with GRCs appearing more active in female oocytes. This restricted but clear transcriptional activity suggests that GRC expression may be developmentally regulated and potentially limited to specific functional roles within the germline, rather than being globally active/repressed. We argue further studies of GRC transcriptional activity would be improved by the use of more sensitive single cell RNA-seq approaches, which would ultimately help elucidate whether GRC-expression is truly as limited to only a small fraction of genes present on these chromosomes.

## Methods

### Fly Husbandry and Strain Used

The *B. coprophila* line used in this study has been maintained as a laboratory stock since the 1920s (Metz 1925). Previous literature has referred to this species as *Sciara coprophila* (Lintner 1895), but it was later reassigned to the genus Bradysia (Steffan 1966). The *B. coprophila* strain, Holo2, was originally obtained from the Sciara stock centre at Brown University in October 2017, and has since been maintained at the University of Edinburgh. The Holo2 strain is descended from the parental line 7298 which carries a dominant phenotypic marker (wavy wing) on the X’ chromosome unique to female-producing females (Metz and Smith 1931) (see ***Supplementary Table S1*** for more information). Eggs of known sex can be collected for *B. coprophila* because mothers are either exclusively male-producing or female-producing (Metz and Smith 1931; Crouse 1960). Fly stocks are maintained by setting up mass cultures of same-sex producing females (i.e., either several gynogenic or androgenic females with a similar number of males per vial) in 28.5mm x 95mm polypropylene vials containing a substrate of 2.2% bacteriological agar. The cultures are kept at 18°C and ∼60% humidity with a 12:12hr light cycle. From the day of hatching until pupation, larvae are fed on a mixture of brewer’s yeast, mushroom powder, spinach powder, nettle powder and ground straw. A small pinch of the mixture is added to each vial twice weekly throughout larval development.

### RNA-Extraction and Experimental Design

We used 42 RNAseq libraries from pooled individuals of both sexes spanning four developmental stages: 0-2 day-old adults, late larvae/early pupa, 4-8h embryos, 0-4h embryos (***Supplementary Table S1***). Adult and pupal/larvae libraries were divided into germline-only (testes or ovaries) and soma-only (remaining carcass) samples. For the adult samples we used publicly-available data from (Baird et al. 2025), which includes 3 male replicates and 6 female replicates (SRA BioProject PRJNA1109384). To generate the pupa/larva libraries, we used larvae in late eyespot stages ranging to early pupa before the eye was more than ¼ full of pigment, as these stages correspond to when meiosis is occurring (just prior to meiosis until the second meiotic division) (Gerbi 2024). We dissected individuals in PBS, using insect pins to detach the germ tissue from the fat body and collecting the germ tissue in one tube and the remaining carcass in another (somatic sample). To collect embryo samples, we mated adult males and females for ∼18-24h, and then pinned females to a petri dish containing 2.2% agar. We applied pressure to the thorax to stimulate egg laying (Gerbi 2024); an entire clutch (50-100 eggs) is usually laid within the following 2 hours. We collected three replicates for each sex for each time stage, with each replicate containing the eggs of approximately 15 females pooled together. We collected embryos at two time stages, one of which precedes GRC elimination from the soma (0-4h), and the other follows GRC elimination (4-8h, de Saint Phalle & Sullivan 1996). We additionally used 3 libraries of each sex from 2h-2 day-old embryos from Urban et al. (2021; BioProject PRJNA291918) to assess GRC transcription during ZGA. To extract RNA, we used a modified version of the PueLink RNA purification kit (ThermoFisher), including a TRIzol solubilisation step. We quantified and quality-assessed samples using a Qubit and Nanodrop (ThermoFisher). We sequenced samples to a depth of approximately 50m reads per library of poly-A selected 150bp paired-end data on the Illumina Novaseq S1 platform.

### RNA-seq

RNA libraries were trimmed to remove adapter sequences using fastp (v0.24.0; Chen et al., 2018) and quality controlled using FastQC (v0.12.1; (Andrews 2012). Paired-end RNA reads were then mapped to the reference *Bradysia coprophila* (GCA_965233685.1; SRA: ERS15411730) genome using STAR (v2.7.11b; Dobin et al., 2013). Annotation for the reference genome was provided to allow STAR to extract splice junctions and improve accuracy of the mapping. This annotation was generated by Hodson et al. (2025) using the EarlGrey TE annotation pipeline (Baril et al. 2024) and two iterative rounds of BRAKER (Gabriel et al. 2024 Feb 29), incorporating both Arthropoda protein data and RNA-seq evidence to refine gene models for core and GRC assemblies. GRC-gene names in the annotation correspond to their sequential position along GRC1 and GRC2, denoted as g1 - g19917, where the numerical suffix indicates the gene’s order along the assembled GRCs. RNA reads were mapped using default parameters, aside from a filter allowing a maximum of two mismatches per paired-end read (--outFilterMismatchNmax 2). This filter reduced mismapping of core chromosome reads to GRC paralogues while avoiding excessive stringency that might discard reads containing sequencing errors or allelic variation. To further reduce any potential mismapping of core genes with GRC paralogues, samtools (v1.21; Li et al., 2009) was utilised to remove any multi-mapped reads. This was achieved by filtering out any reads that did not have a mapping quality flagged as 255, which STAR assigns to any reads mapping uniquely to the reference genome. Transcript per million (TPM) normalisation was performed on uniquely mapped alignment files using StringTie (v2.2.3; Kovaka et al., 2019), to account for both sequencing depth and transcript length.

### Defining Expression Threshold for Expression Using Intergenic Region Mapping

For this study, we define a gene as expressed if it has a TPM that is statistically significant above the background mismapping rate in at least 2 of the three replicates. To quantify the rate of mismapping as background noise in our RNA-seq data, we estimated TPM of reads mapping to presumed inactive intergenic regions, following a method proposed by (Costa et al. 2022). Similar to their approach, we defined intergenic regions as genomic intervals located at least 500 bases from any annotated gene, and of a minimum length of 1 kb. For any intergenic regions exceeding 20 kb in length, only the central 20 kb segment was retained to avoid inclusion of potentially unannotated features at the margins. We implemented a custom Python script to generate an intergenic-only GTF file from our genome assembly and corresponding gene annotation. Uniquely mapped reads for all life stages for each species were then re-quantified using StringTie (v2.2.3, Kovaka et al., 2019), with the intergenic GTF provided as input, yielding TPM values for each intergenic region. Due to imperfections in gene annotations common in non-model species, we expected that some actively transcribed genes or non-coding RNAs might remain within the defined intergenic regions. Therefore, to more accurately characterise the background mismapping to non-functional intergenic regions of the genome, we adopted a deconvolution-based method to separate background signals from genuinely expressed elements (Costa et al. 2022). Specifically, we applied a Gaussian Mixture Model (GMM) using the Mclust package in R (Scrucca et al. 2016) to the TPM values of intergenic regions. The optimal number of components was selected based on the Bayesian Information Criterion, and each region was assigned to a component. We designated the component with the lowest overlap with the curve representing the expression of annotated coding regions as representing the inactive, background-like intergenic signal as suggested in Costa et al., (2022) (***Supplementary Figure S1***). The maximum TPM observed in this component was retained as the MITT_species_ (Maximum Inactively Transcribed Threshold), serving as a species-wide expression threshold to define background transcription. All TPMs below the MITT_species_ value were selected to form a reference intergenic set.

Using TPM values of the reference intergenic set, we calculated the mean and standard deviation. These statistics were then used to compute a Z-score for each *B coprophila* gene with associated expression (global across libraries). One-tailed p-values were derived from the standard normal distribution, representing the probability of a gene’s expression being due to background transcription alone. To control for multiple hypothesis testing, p-values were adjusted using the Benjamini-Hochberg procedure (Benjamini and Hochberg 1995). The smallest TPM value for which the adjusted p-value was less than or equal to 0.05 was taken as the MATT_library_ (Maximum Actively Transcribed Threshold) threshold. Genes with TPM values above this cutoff were classified as “active”. Distributions of TPM values were compared between intergenic and genic datasets, and all visualisation (***Supplementary Figure S1***) was performed using ggplot2 (Valero-Mora 2010) and patchwork (Pedersen 2025).

### Filtering of Confidently GRC-expressed Genes

Genes were considered to be expressed only if they surpassed the active TPM threshold (∼0.22) in at least two libraries per stage. Erroneous mapping has been noted in transcriptomic analysis of GRCs, likely due to the presence of highly similar core chromosome paralogues on the GRCs (Schlebusch et al. 2023) or the contamination of somatic libraries with germline tissue. In the case of the former, the issue emerges when reads originating from core loci may align to paralogous GRC-linked genes, producing false-positive signals of GRC expression. Crucially, such mismapping has been shown to persist even when filtering for uniquely mapped reads (Schlebusch et al., 2023). To reduce false positives from core chromosome mismapping, GRC-linked genes expressed in germline libraries were excluded if they also showed expression above the TPM threshold in matched somatic libraries (**Figure 1B,C**). Because GRCs are absent from somatic cells, any apparent expression in these libraries may reflect misaligned core-derived reads. These same reads could also falsely elevate expression estimates in germline libraries, making such GRC genes unreliable for this analysis. Additionally, genes with significant BLAST similarity to core genome sequences were not considered confidently GRC-expressed (defined as having %identity x %coverage over 70), as their paralogous relationships may further confound expression. This BLAST-based filtering was applied across all stages, including embryos, which lacked somatic control libraries. Finally, read pile-ups were manually inspected to confirm that reads mapped unambiguously to GRC loci (***Supplementary Figure S3***).

Whilst we did not specifically correct for germline contamination, we did investigate the expression of four canonical germline genes in male and female germline versus somatic libraries (***Supplementary Figure S2***). This was achieved by first identifying homologues of these genes in *B. coprophila* core genome using tblastx and the flybase CDS sequences (Altschul et al. 1990; Camacho et al. 2009). TPM values for these genes in the different libraries were generated from the aforementioned RNA-seq data generated in this study.

### Homology Analysis for Expressed GRC-linked genes

To investigate the potential functions of the 15 GRC-linked genes confidently expressed across *B. coprophila* development, homology-based analyses at the protein level were performed. Protein sequences of GRC-linked genes were queried against the NCBI non-redundant (nr) protein database using BLASTp (v2.16.0) to identify similarities to known proteins (Altschul et al. 1990). In parallel, Repbase was searched against to assess homology with transposable elements and other repetitive DNA sequences (Bao et al. 2015) (**Figure 5A**). Additionally, all proteins were queried through InterProScan (v5.76-107.0; (Zdobnov and Apweiler 2001) in order to infer function through protein family classification and domain prediction.

### Identifying HGT Candidate Using Long Read Data

We re-mapped the PacBio long-read sequencing data, originally used to assemble the *B. coprophila* reference genome (GCA_965233685.1), back to the annotated and assembled genome using minimap2 (2.28-r1209; Li, 2018). A custom python script was used to calculate GC content across the putative HGT region, using a 2kb sliding window sliding at a step of 1kb. Figures were generated using IGV, with minor indels (>100bp) hidden for viewing clarity. Alignments between *B coprophila* GRC2 and *Rickettsia* were performed using FASTGA (v1.1; https://github.com/thegenemyers/FASTGA) with default parameters. The alignment was visualised using a dot plot generated using a custom R script.

### Cytological Assays to Identify GRC Transcription

Immunostaining experiments were performed as previously described in (Cheng et al. 2008) with minor modifications. Briefly, gonads from 10 to 15 individuals at the desired stage (late larvae or adults) were dissected in 1X PBS, fixed in 4% formaldehyde (Thermo Scientific, Cat#10751395) for 30min at room temperature, then washed three times for 30min in 0.1% PBS-T (1X PBS, 0.1% Triton (Merck, Cat#X100). After washes, samples were permeabilised in 0.3% PBS-T for 1h30min at room temperature, followed by a brief wash in 0.1% PBS-T, then incubated with primary antibody in 0.1% PBS-T at 4°C overnight. Samples were washed three times in 0.1% PBS-T for 20min, then incubated with secondary antibody and with 1µg/ml Hoechst stain (biorbyt Cat#orb363806) in 0.1% PBS-T at 4°C overnight, washed again three times for 20min, and mounted in mounting medium (80% Glycerol (Merck, Cat#G-5516), 1X PBS, 0.2% propyl gallate (Merck, Cat#P3130). Primary antibodies used were: rabbit anti-H4K16ac (1:100, Active Motif Cat# 39929, RRID:AB_2753164); rat anti-Vasa (1:100, developed by Spradling, A.C. and Williams, D., DSHB Cat# anti-vasa, RRID:AB_760351). Secondary antibodies used were: Goat anti-rabbit IgG Alexa488 (1:300, Abcam Cat#ab150077, RRID:AB_2630356), Goat anti-rat IgG Alexa555 (1:300, Thermo Fisher Scientific Cat#A-21434, RRID:AB_2535855).

### Image Acquisition and Processing

Immunofluorescence images were acquired on a Leica SP5 confocal microscope with 40x oil-immersion objective and processed with Leica’s LAS X (RRID:SCR_013673) and Fiji (RRID: SCR_002285) softwares.

## Data availability

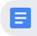 [Supplementary_Materials](https://docs.google.com/document/d/1D4hNGQOPKm19O8s4FHTI92jYcS20MsosfXuDkdF2E00/edit?tab=t.0#heading=h.2h3t6tifta47)

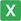 [Supplimentary_data.xlsx](https://docs.google.com/spreadsheets/d/1ZjjYUNeQekw4x_c-9vi2csk_CyhzUqt0/edit?gid=485512448#gid=485512448)

Code repository: https://github.com/RiccardoKyriacou/GRC_transcription

Reference *Bradysia coprophila* genome: GCA_965233685.1; SRA: ERS15411730

The embryo RNAseq data is available on SRA under BioProject accession PRJNA1220056.

The pupa/larval RNAseq data and *Rickettsia* genome will become available upon publication

## Acknowledgements

We would like to thank the Ross lab, Kamil Jaron, the Tree of Life Program at Wellcome Sanger Institute, and Judith Mank for helpful comments during the preparation of this manuscript. This research was funded though a Royal Society Newton International Fellowship (NIF\R1\232397, C.N.H), a European Research Council grant (PGErepro, grant agreement 803932, L.R) and a Royal Society Dorothy Hodgkin Fellowship (DHF\R1\180120, L.R and R.G.K)

