## Supplementary material for "Evidence for Transcription and Horizontal Gene Transfer in Dipteran Germline-Restricted Chromosomes": Supplementary_materials.pdf

#### Supplementary Figures

Supplementary Figure S1: Defining Active Expression Threshold for GRC-linked Genes

Supplementary Figure S2: Expression of known germline-specific genes in somatic versus germline libraries

Supplementary Figure S3: RNA read pileups for 15 confidently expressed GRC-linked genes

Supplementary Figure S4: Sex differences in expression for confidently expressed GRC-linked genes

Supplementary Figure S5: Read Counts for DNA-reads Spanning Across HGT-region Boundary

Supplementary Figure S6: Example of bacterial RNA mapping to a GRC-linked gene of bacterial origin (g19121) within HGT-region of GRC2

#### Supplementary Tables

Supplementary Table S1: RNA-seq library breakdown

Supplementary Table S1: Pooled Embryo RNA-seq Mapping

Supplementary Table S3: Summary BLAST results for expressed GRC-linked genes against *L ingenua* and *B. impatiens* genome

### Supplementary Figures

#### Supplementary Figure S1: Defining Active Expression Threshold for GRC-linked Genes

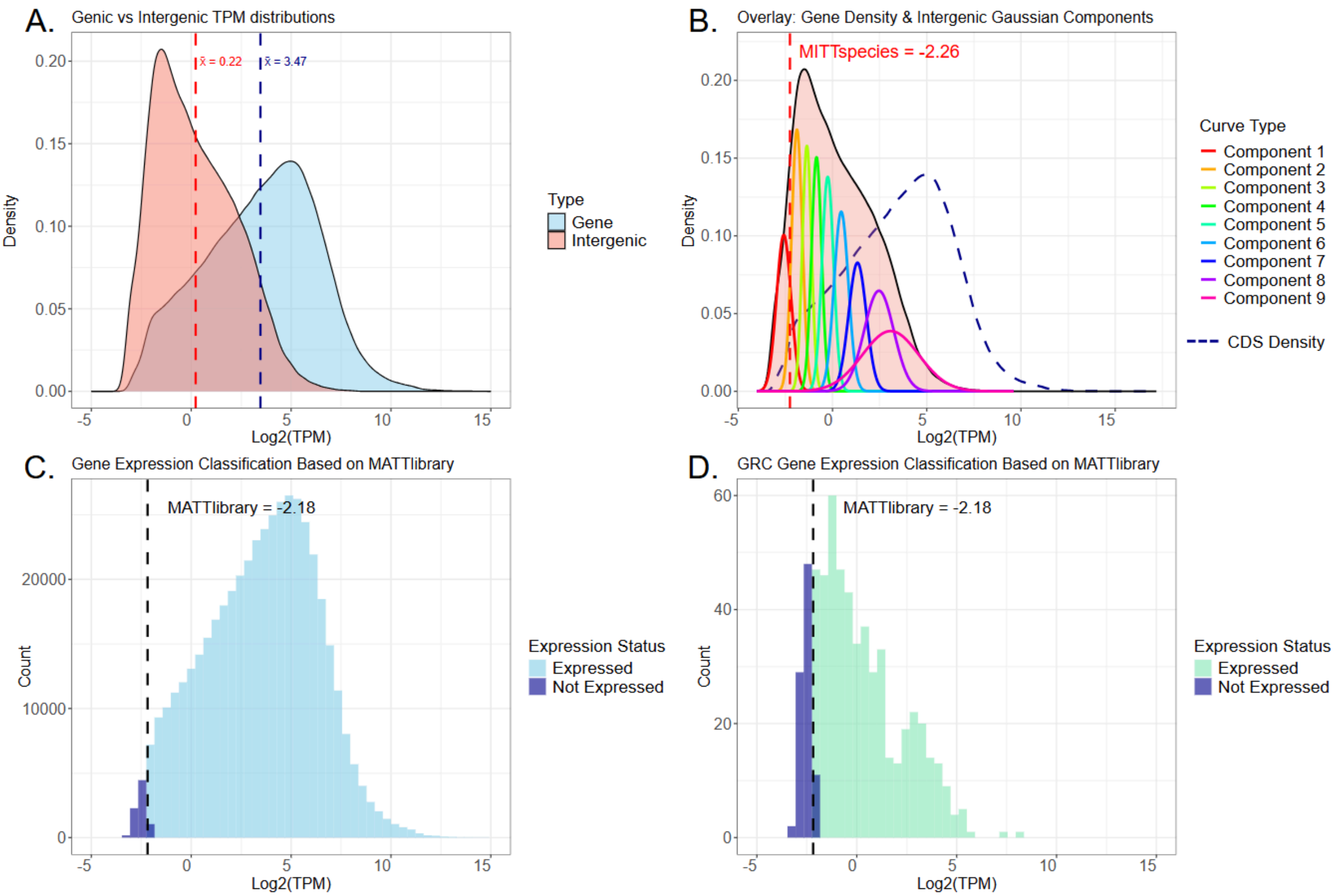


**Fig. S1: Approach to define active transcripts-per-million (TPM) threshold for GRC-linked genes using intergenic region mapping rate (Costa et al., 2022)**

**A.** Log2(TPM) density curves for genes and intergenic regions. Intergenic regions are ≥500 bp from any annotated gene and ≥1 kb long. Mean intergenic = 0.22 log2(TPM) (1.2 TPM), mean coding genes = 3.47 log2(TPM) (11.1 TPM). Given imperfect annotation in the non-model species *Braydesia coprophila*, some unannotated genes, active TEs, and other RNAs are expected within intergenic space.

**B.** The intergenic log2(TPM) distribution was deconvolved with a Gaussian mixture (mclust), with the number of components selected by BIC (here, K = 9). Overlaid is the coding sequence (CDS) gene density (dark blue, dashed). By visual inspection, we selected the component with minimal apparent overlap with the gene density to represent background transcription, as per Costa et al., (2022). We then defined a species-level upper bound for background (MITTspecies, red vertical dotted line) from this component and used it as a threshold.

**C.** Using intergenic regions with log2(TPM) below MITTspecies as a reference background set, we estimated a global (pooled across libraries) background mean and standard deviation. For each gene, we computed one-tailed p-values under this background model and adjusted them with the Benjamini–Hochberg procedure. The smallest TPM at which the adjusted p-value ≤ 0.05 defines the library-wide MATTlibrary (vertical black line, dashed) threshold. MATTlibrary = -2.18 log2(TPM) (~0.22 TPM). Genes above MATTlibrary are classified as “active” (light blue); genes below are “inactive” (dark blue).

**D.** Gene expression on the GRC scaffolds. The dashed vertical line indicates the MATTlibrary threshold from **C.** Genes above the threshold are “active” (green) and those below are “inactive” (dark blue).

#### Supplementary Figure S2: Expression of known germline-specific genes in somatic versus germline libraries

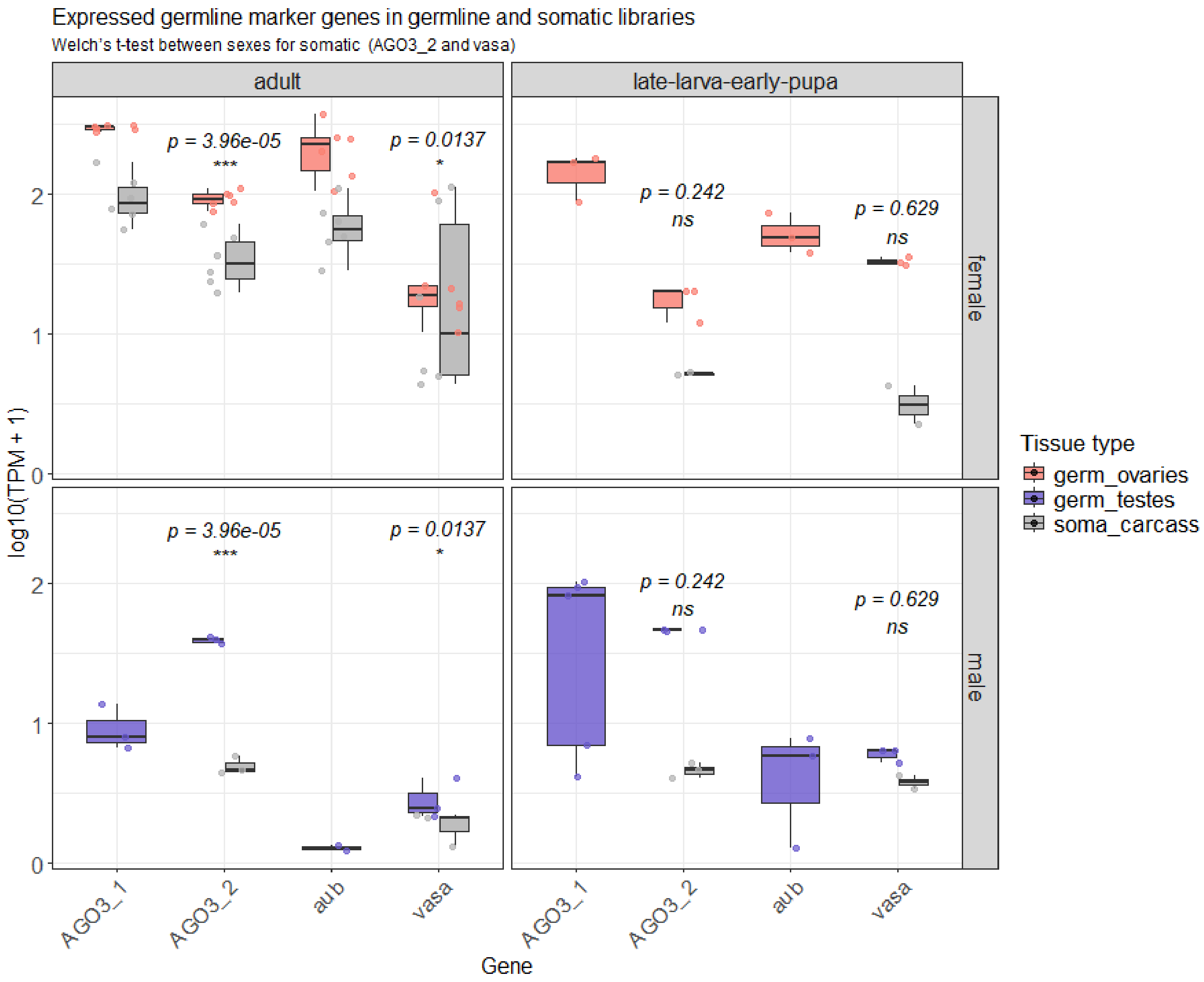


**Fig. S2: Expression of germline marker genes in *B. coprophila* somatic versus germline tissues across sexes and developmental stages.** Log-transformed TPM values log10(TPM + 1) are shown for four germline-specific genes: *aubergine* (*aub*), *vasa*, and two *Argonaute 3* homologues (*AGO3 1* and *AGO3 2*). TPM values are plotted for germline libraries (testes, blue; ovaries, pink) and somatic carcass libraries (grey). To ensure robust comparisons, only genes with TPM > 0.22 in ≥2 libraries per stage/tissue were retained (see ***Fig. S1***). Data are faceted by sex (rows: female, male) and developmental stage (columns: late-larva–early-pupa, adult).

*AGO3 1* and *aubergine* only show somatic expression (grey boxes) in female adult libraries. *AGO3 2* and *vasa* show somatic expression in female and male somatic libraries in both life stages. For genes showing germline contamination in both sexes (*AGO3 2* and *vasa),* differences in TPMs in the somatic carcass libraries were formally tested using Welch's t-test. Significance is annotated above the boxes: *** $p < 0.001$, ** $p < 0.01$, * $p < 0.05$, ns = not significant. Results indicate female-biased somatic contamination in both *AGO3 2* and *vasa* in adults, whereas late-larval/early-pupal stages show no significant sex differences.

*Shapiro-Wilk normality test for log10(TPM + 1):*

| Gene | Stage | Sex | n | p-value |
|---|---|---|---|---|
| AGO3_2 | adult | female | 6 | 0.841 |
| AGO3_2 | adult | male | 3 | 0.300 |
| AGO3_2 | late-larva-early-pupa | male | 3 | 0.942 |
| vasa | adult | female | 6 | 0.100 |
| vasa | adult | male | 3 | 0.091 |

*Levene's test for equality of variance:*

| Gene | Stage | p-value |
|---|---|---|
| AGO3_2 | adult | 0.0829 |
| AGO3_2 | late-larva-early-pupa | 0.328 |
| vasa | adult | 0.0758 |
| vasa | late-larva-early-pupa | 5.55e-32 |

*Welch's t-test comparing log10(TPM + 1) between sexes (somatic carcass libraries):*

| Gene | Stage | p-value | Female mean | Male mean | Difference (F–M) |
|---|---|---|---|---|---|
| AGO3_2 | adult | 3.96e-05 | 1.52 | 0.696 | 0.826 |
| AGO3_2 | late-larva-early-pupa | 0.242 | 0.716 | 0.665 | 0.051 |
| vasa | adult | 0.0137 | 1.22 | 0.265 | 0.959 |
| vasa | late-larva-early-pupa | 0.629 | 0.493 | 0.581 | –0.089 |

#### Supplementary Figure S3: RNA read pileups for 15 confidently expressed GRC-linked genes

##### g233 - late larval/early pupa

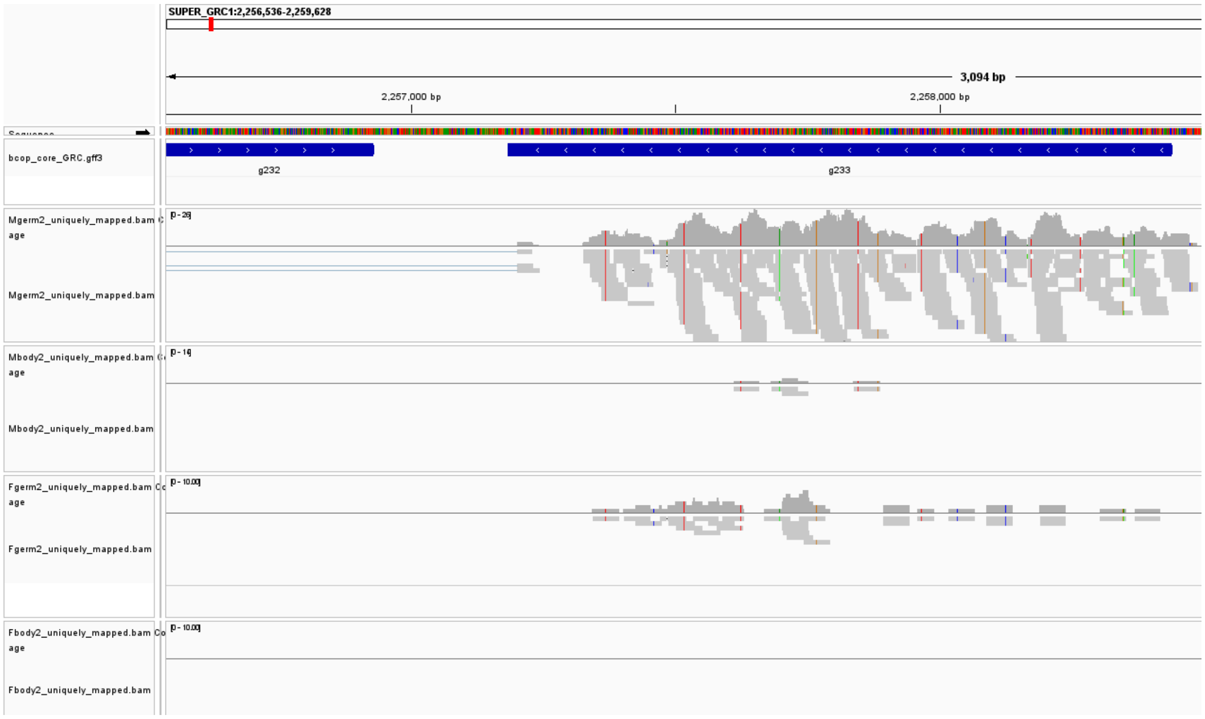


##### g491 - late larval/early pupa

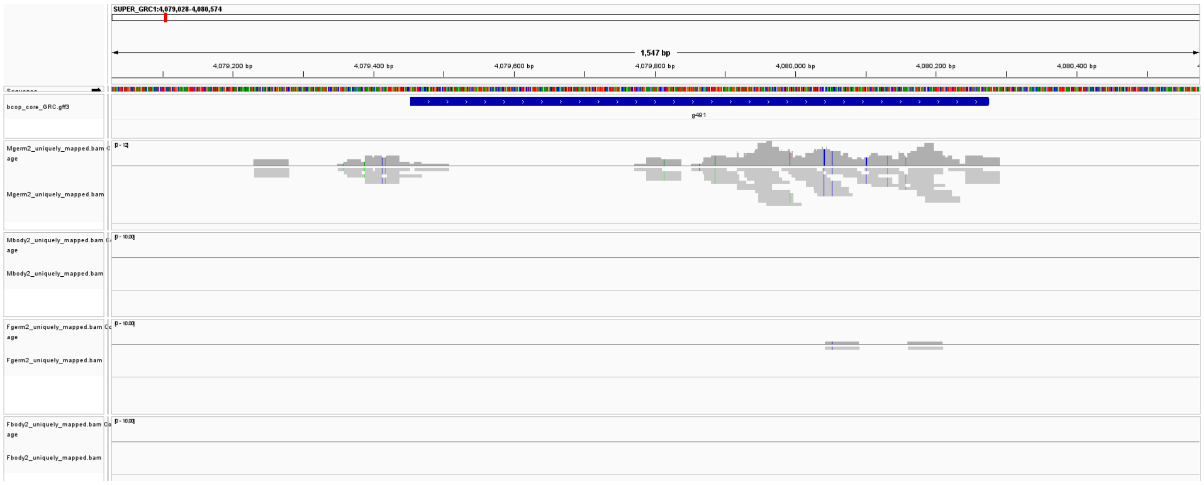

### g596 - late larval/early pupa

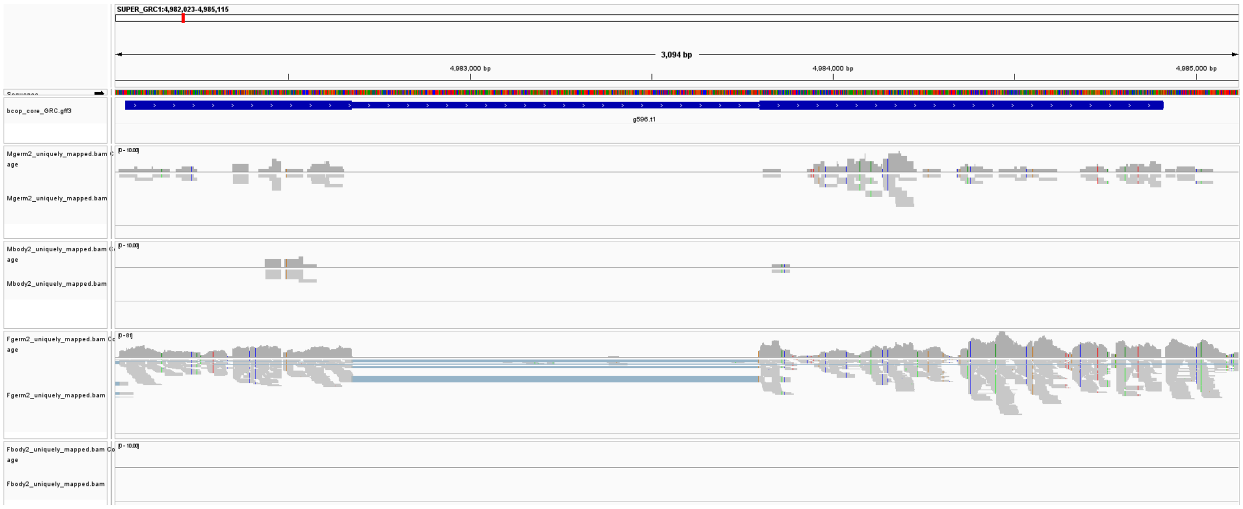


### g7610 - adult

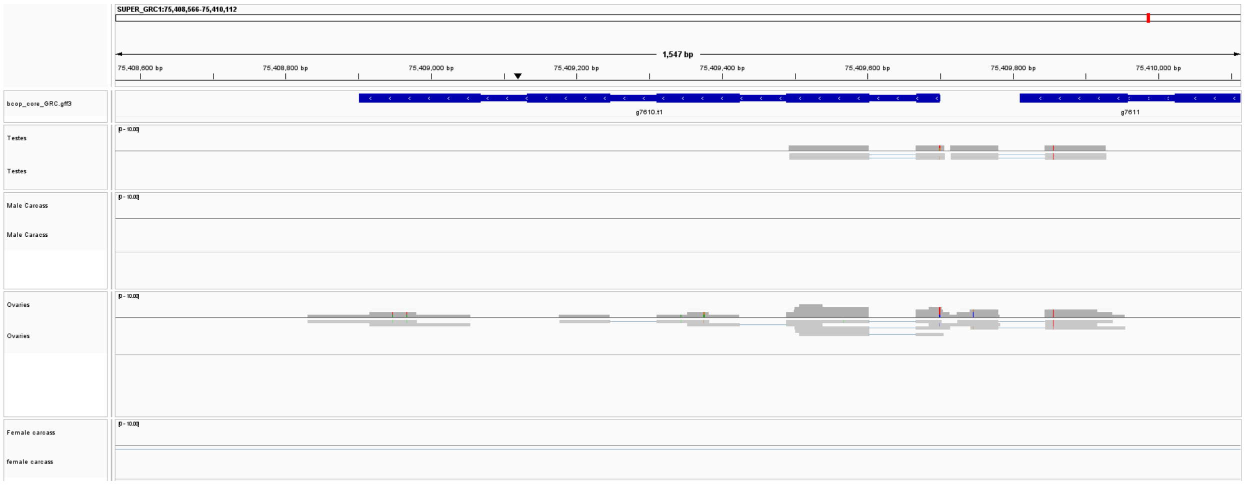


### g7957 - 4-8h embryo

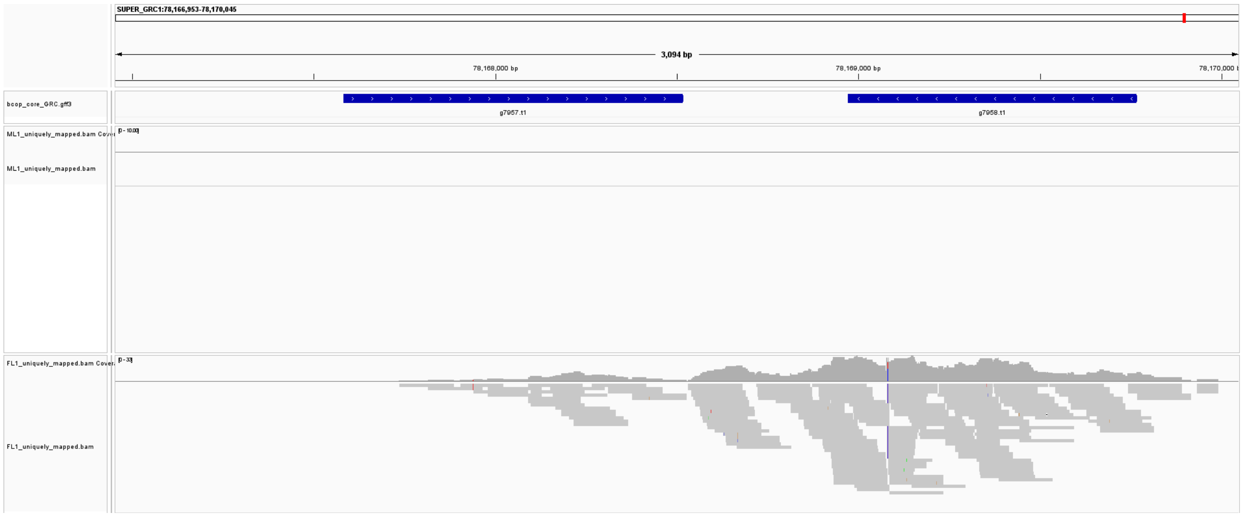

### g8036 - 4-8h embryo

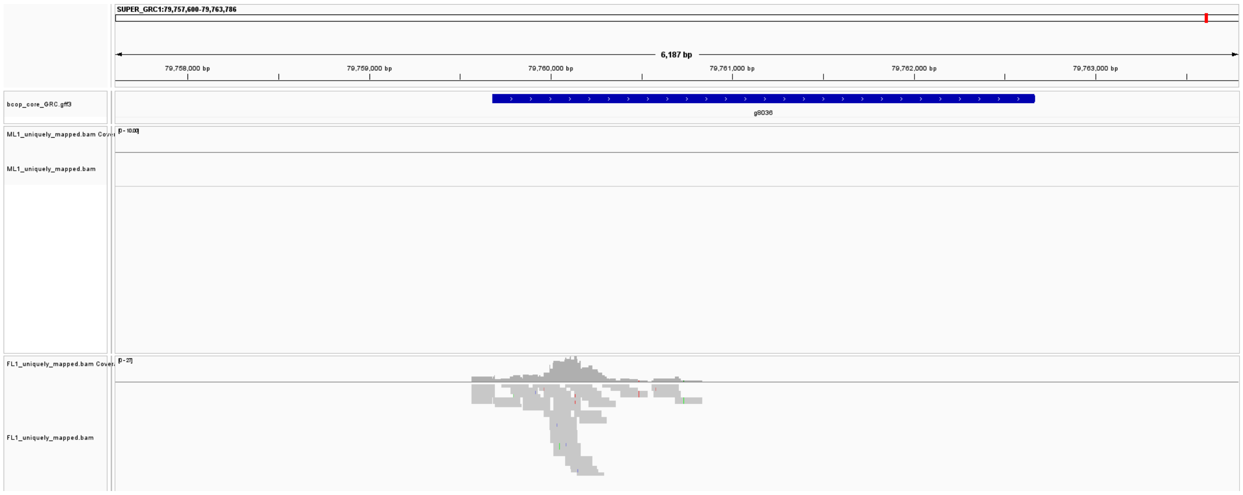


### g11713 - adult

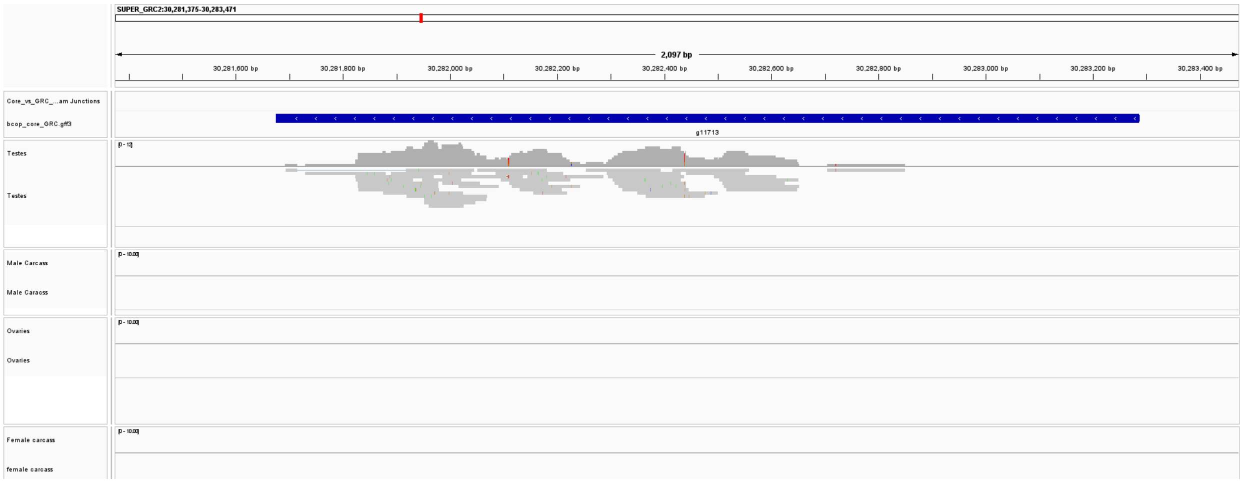


### g13362 - adult

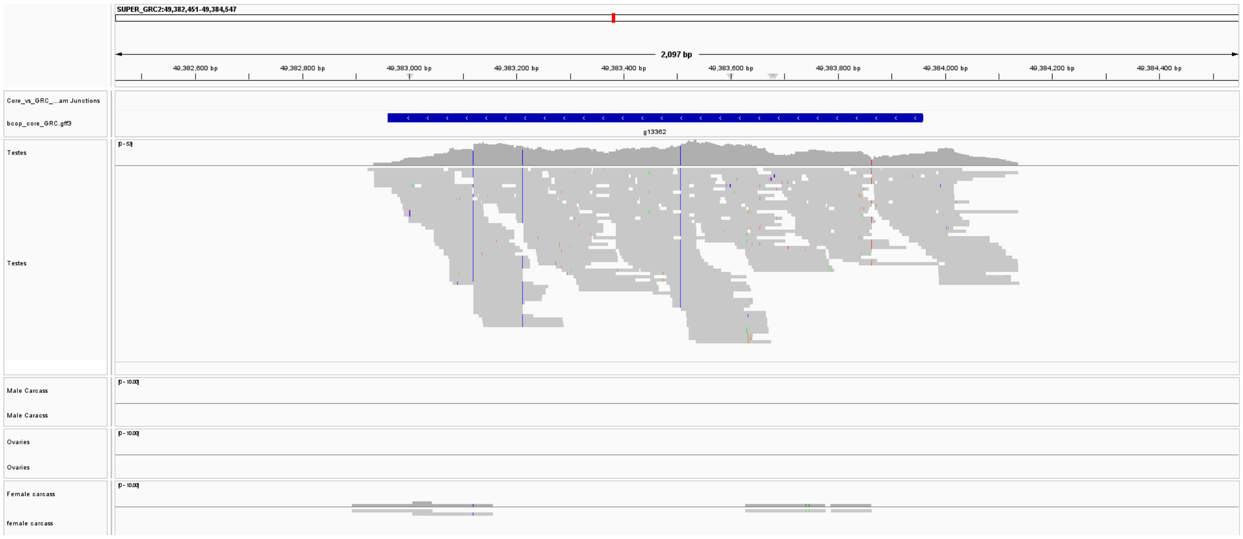

#### g13362 - late larval/early pupa

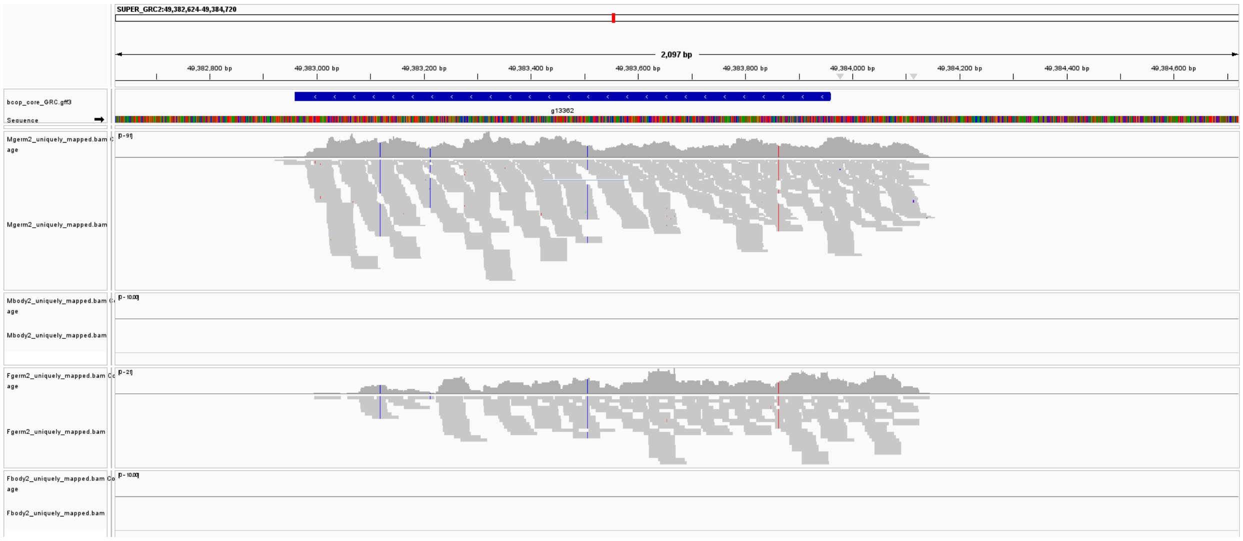


#### g13694 - adult

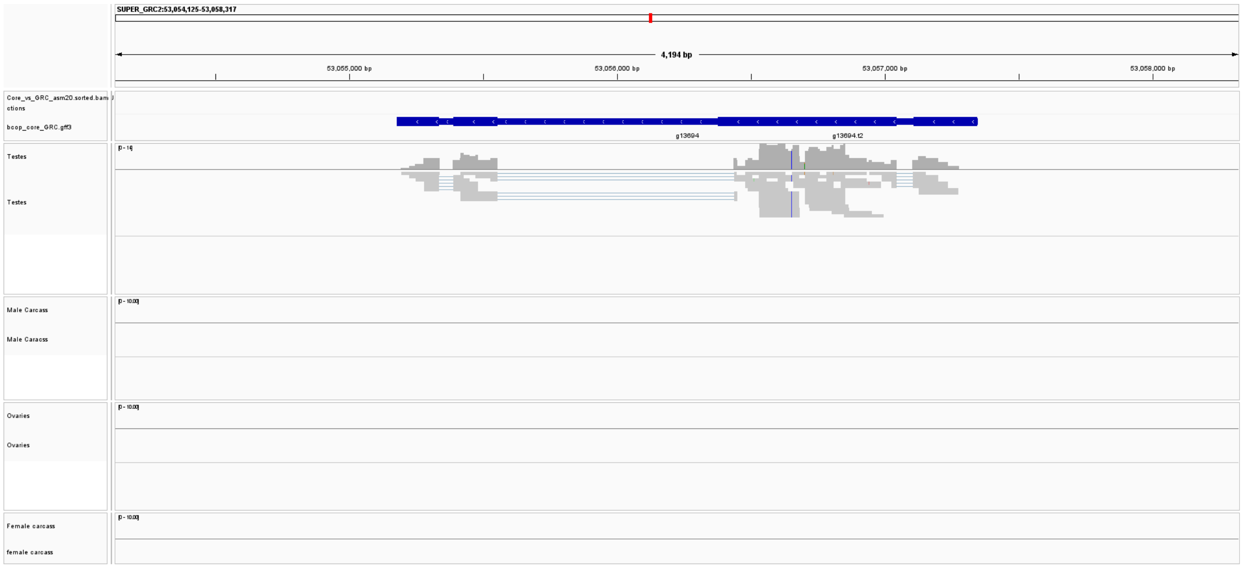


#### g13694 - late larval/early pupa

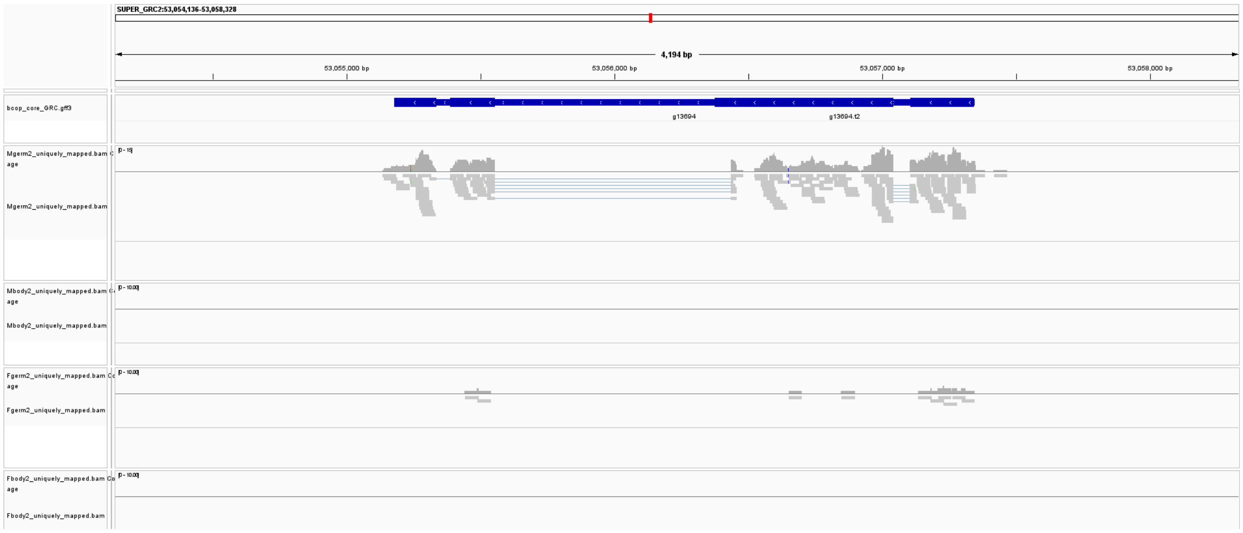

#### g15174 - adult

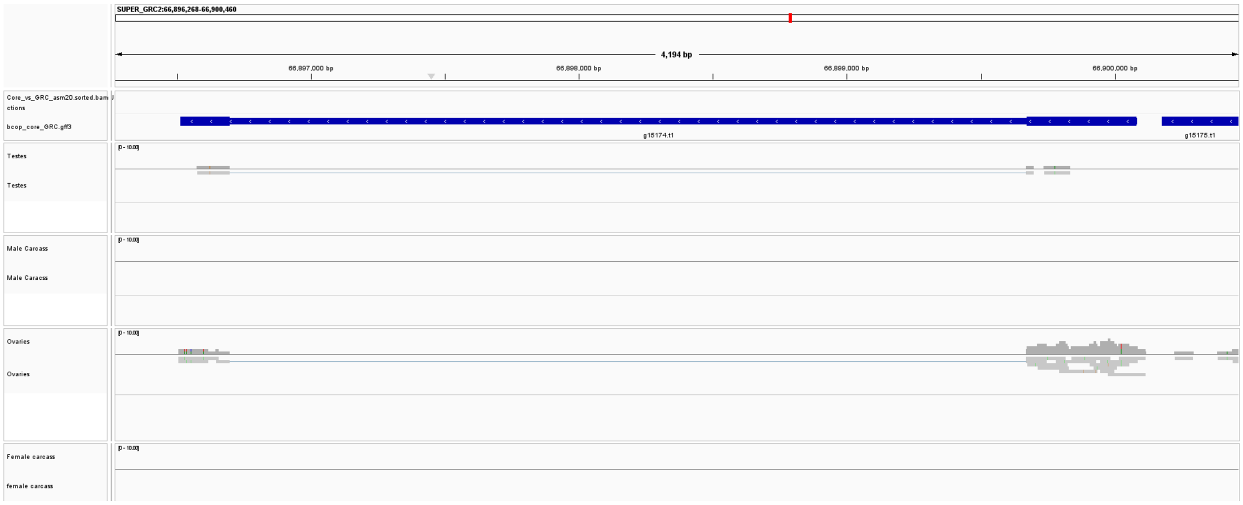


#### g16029 - late larval/early pupa

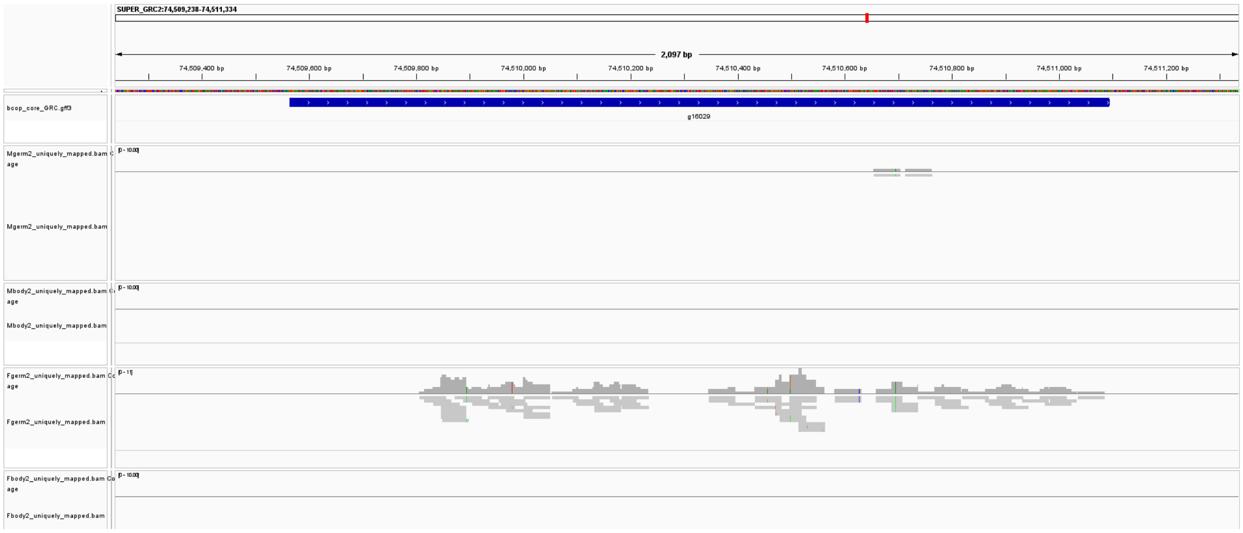


#### g17107 - adult

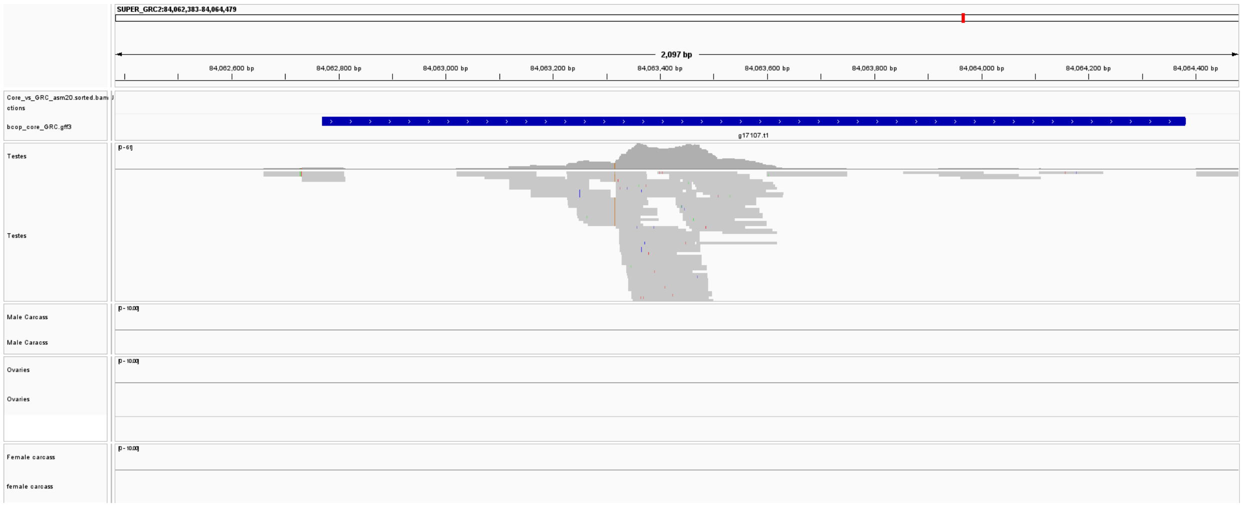

#### g17119 - adult

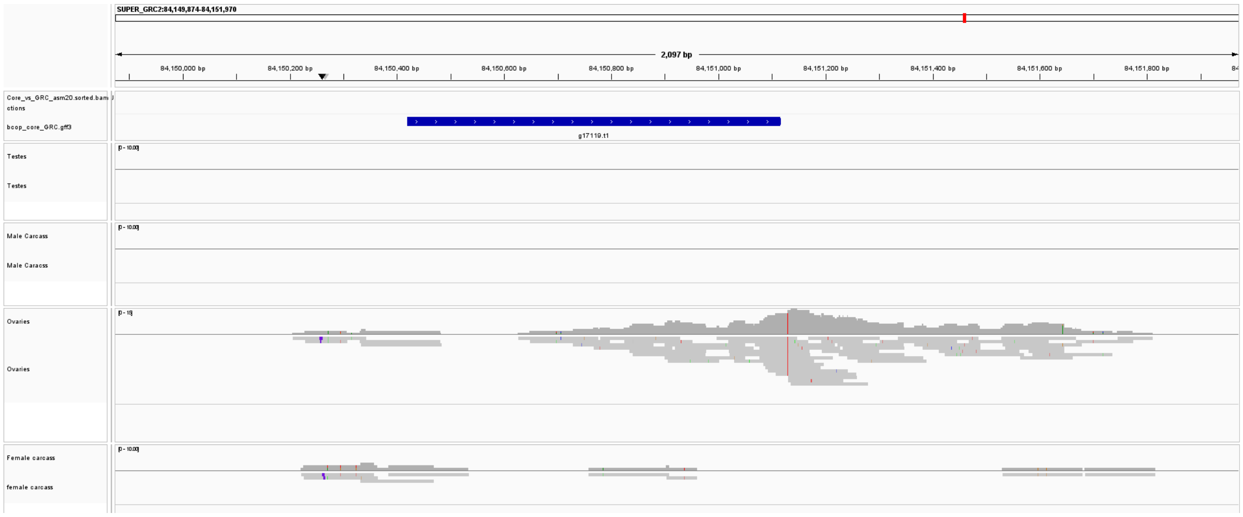


#### g19161 - late larval/early pupa

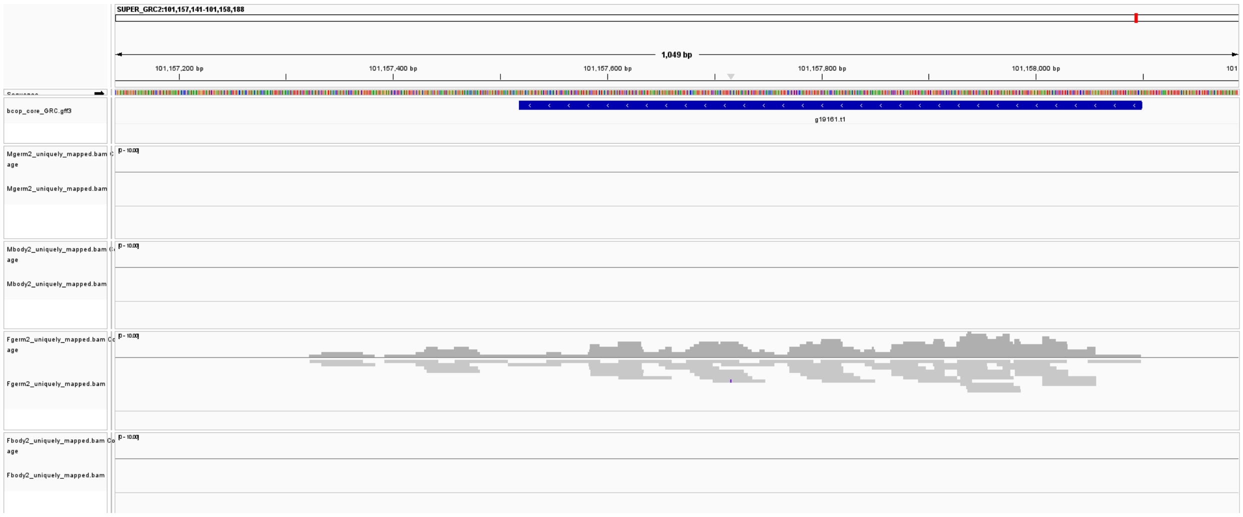


**Fig. S3: Integrated Genomics Viewer ( IGV) Robinson et al. (2011) screenshots showing RNA read pile ups for 15 GRC-linked GRC genes identified in this study.** For genes expressed at the 0-4h and 4-8h embryo stage one male library and one female library are shown. For late larvae/early pupae and adult, matched carcass and germline libraries are shown for male and female.

#### Supplementary Figure S4: Sex differences in expression for confidently expressed GRC-linked genes

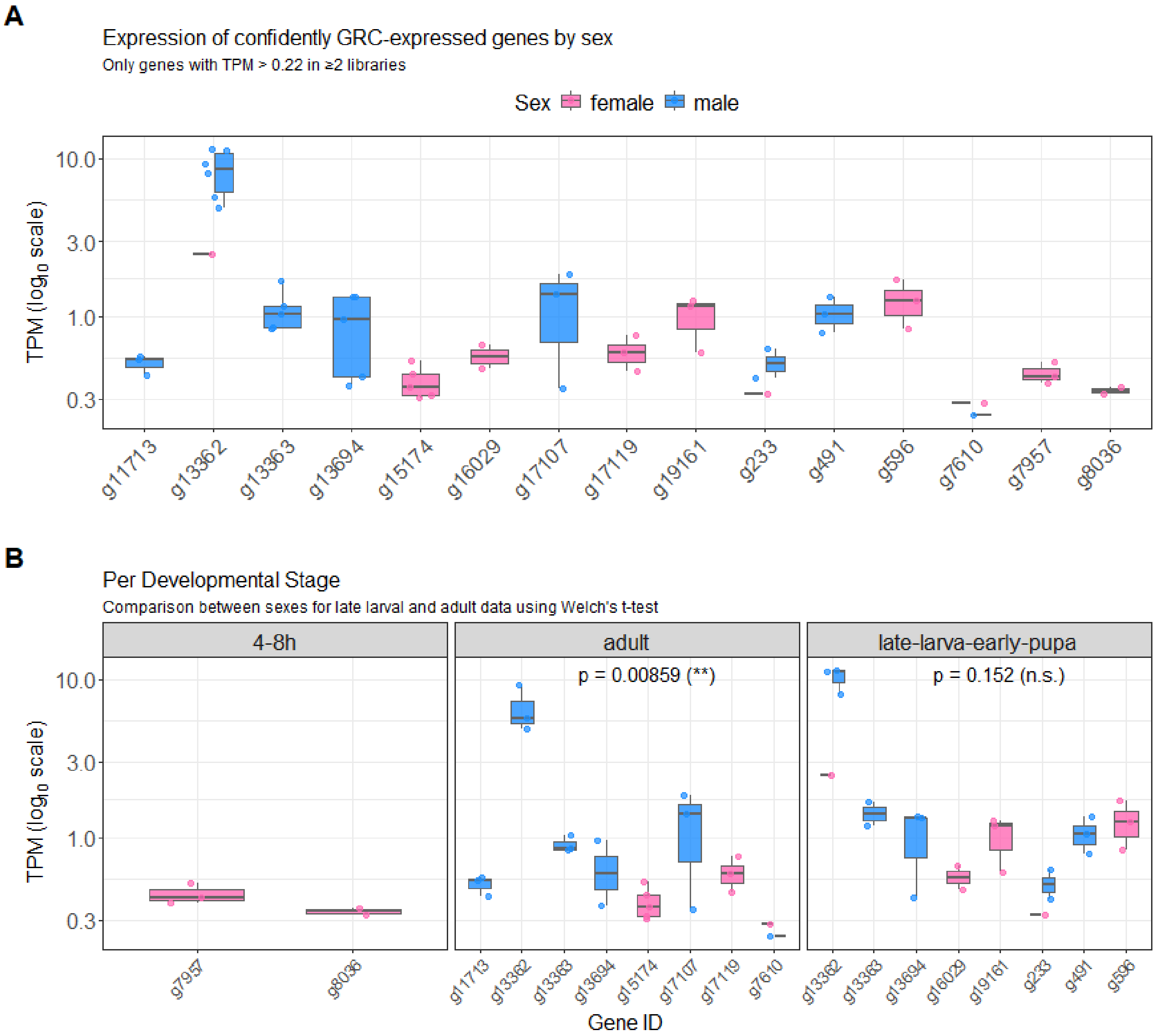


**Fig. S4: Boxplots showing the expression (TPM, log10-transformed) of confidently GRC-expressed genes across developmental stages in males and females.** Only genes with TPM > 0.22 in ≥2 libraries were included, with no similar core chromosome paralogues or somatic mismapping (see Main Methods).

**A.** Plot for all stages combined. Individual data points are overlaid as jittered points.

**B.** Plot faceted by developmental stage. Statistical comparisons between sexes were performed using Welch's t-test for each developmental stage, with p-values and significance annotations shown above each facet (* $p < 0.05$, ** $p < 0.01$, *** $p < 0.001$, n.s. = not significant). Adult males exhibit significantly higher expression of GRC-expressed genes compared to females ($p = 0.0086$), whereas no significant difference is observed at the late-larva-early-pupa stage ($p = 0.152$). 4-8h data had no male comparisons.

*Shapiro–Wilk test for normality of log10(TPM) per sex and stage:*

| Stage | Sex | n | Shapiro-Wilk p-value |
|---|---|---|---|
| 4–8h | female | 5 | 0.768 |
| adult | female | 9 | 0.755 |
| adult | male | 15 | 0.210 |
| late-larva-early-pupa | female | 10 | 0.974 |
| late-larva-early-pupa | male | 13 | 0.0327 |

*Levene's test for homogeneity of variance between sexes per stage (note: 4–8h stage only had female samples hence Levene's and Welch test not applicable):*

| Stage | Levene p-value |
|---|---|
| adult | 0.0358 |
| late-larva-early-pupa | 0.301 |

*Welch's t-test comparing log10(TPM) between sexes per stage*

| Stage | Female mean | Male mean | Difference (F − M) | p-value | Significance |
|---|---|---|---|---|---|
| adult | −0.373 | 0.015 | −0.388 | 0.0086 | ** |
| late-larva-early-pupa | −0.041 | 0.201 | −0.242 | 0.152 | n.s. |

#### Supplementary Figure S5: Read Counts for DNA-reads Spanning Across HGT-region Boundary

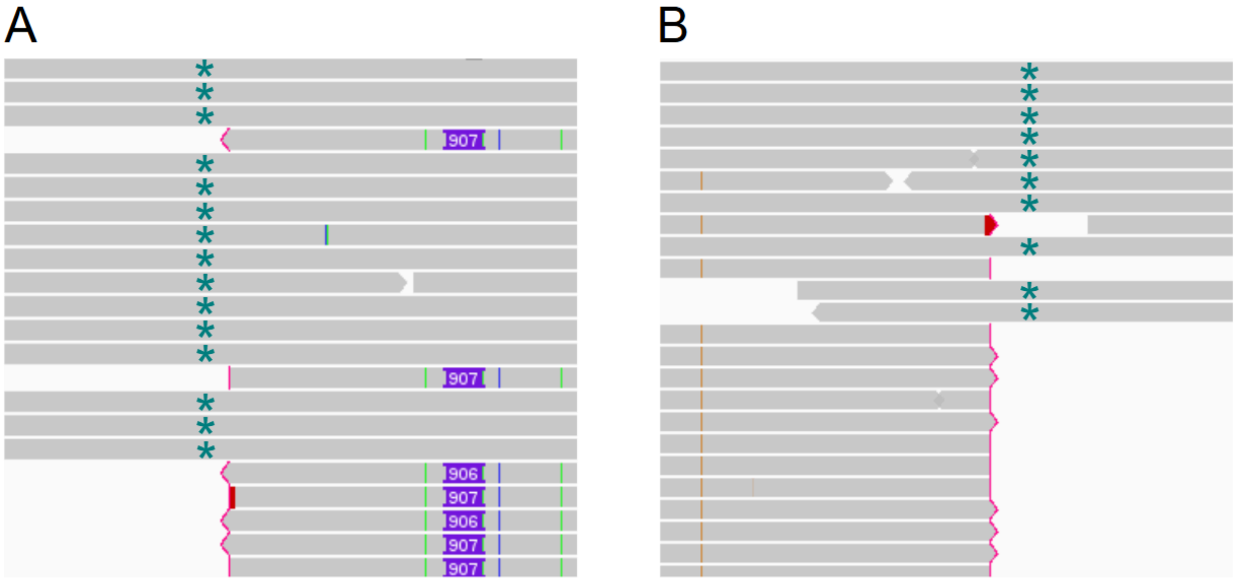


**Fig. S5: IGV visualisation of PacBio HiFi long-read alignments spanning the putative horizontal gene transfer (HGT) region on GRC2.** Shown are individual HiFi DNA reads mapped across the boundary of the putative HGT region on the GRC2 scaffold (green asterix). These alignments demonstrate that the HGT region is physically integrated into GRC2. Reads terminating at the edge of the HGT region (soft-clipped reads, indicated by red vertical lines) represent bacterial-derived sequences that map exclusively within the HGT region but not to adjacent GRC2 sequence.

**A.** 15 long reads (dark green asterix) span across the high coverage/HGT region on its leftmost boundary (downstream). All 15 of these reads do not have diagnostic SNPs/indels shared with bacterial-derived DNA reads. Green SNP: G → A, blue SNP: T → C

**B.** 10 reads (dark green asterix) span across the high coverage/HGT region on its rightmost boundary (upstream). All 10 of these reads do not have diagnostic SNPs shared with bacterial-derived DNA reads. Brown SNP: A → G

#### Supplementary Figure S6: Example of bacterial RNA mapping to a GRC-linked gene of bacterial origin (g19121) within HGT-region of GRC2

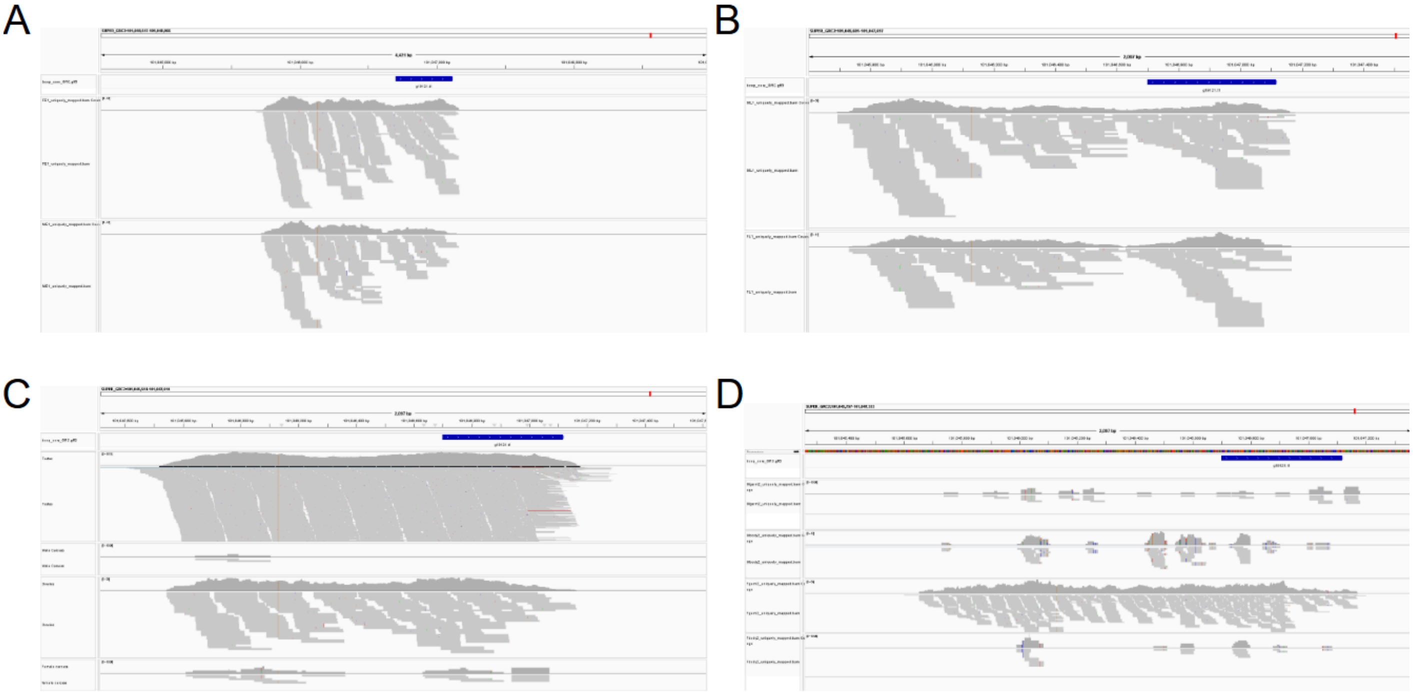

**Fig. S6: IGV screenshots showing RNA-read pileups for GRC-linked gene *g19121*.** This gene is located in the putative HGT region of GRC2 (see Main Results), hence shares extremely close sequence similarity to a gene in the genome of the *Rickketsia* endosymbiont donor. *g19121* shows extremely high TPM and read count (compared to other GRC genes) across all developmental life-stages, and thus is likely to be an artefact of bacterial RNA (present in the germline libraries) mapping to GRC2. For this reason, *g19121* was excluded from our confidently expressed GRC-linked gene dataset and downstream analyses.

**A.** RNA reads mapping to GRC-linked gene g19121 in 0-4h embryo female (top) and male (bottom) libraries.

**B.** RNA reads mapping to GRC-linked gene g19121 in 4-8h embryo male (top) and female (bottom) libraries.

**C.** RNA reads mapping to GRC-linked gene g19121 in late larval/early pupal libraries in males (testes above, carcass below) and females (ovaries above, carcass bottom).

**D.** RNA reads mapping to GRC-linked gene g19121 in adult libraries in males (testes above, carcass below) and females (ovaries above, carcass bottom).

### Supplementary Tables

#### Supplementary Table S1: RNA-seq library breakdown

| **Library description** | **Development Stage** | | | |
|---|---|---|---|---|
| | **0-4h** | **4-8h** | **late larva/early pupa** | **adult** |
| **Male GRC-containing (pre-GRC elimination)** | 3 | 3 | | |
| **Female GRC-containing (pre-GRC elimination)** | 3 | 3 | | |
| **Male GRC-containing (testes)** | | | 3 | 3 |
| **Male GRC-negative (carcass)** | | | 3 | 3 |
| **Female GRC-containing (ovaries/eggs)** | | | 3 | 6 |
| **Female GRC-negative (carcass)** | | | 3 | 6 |

**Total RNA libraries: 42**
**Reference genome: *Bradysia coprophila* (GCA_965233685.1)**
**https://www.ncbi.nlm.nih.gov/datasets/genome/GCA_965233685.1**

*Female libraries contained 3 replicates for both androgenic (male producing) and gynogenic (female only producing) females, hence the resulting six libraries. *B. coprophila* is a monogenic species whereby females will produce either only sons or only daughters. This monogeny is genetically determined in *B. coprophila* due to an inversion on the X chromosome (X'). X' females (female producing) are identified by a phenotypic marker (wavy wing) on the X' chromosome unique to gynogenic females (Metz and Smith, 1931).

#### Supplementary Table S2: Pooled Embryo RNA-seq Mapping

| GRC Gene ID | GRC Scaffold | Average TPM across libraries |
|---|---|---|
| **g6396** | SUPER_GRC1 | 11.73 |
| **g13065** | SUPER_GRC2 | 5.06 |
| **g14257** | SUPER_GRC2 | 3.00 |
| **g14637** | SUPER_GRC2 | 15.60 |
| **g16029*** | SUPER_GRC2 | 5.87 |
| **g17249** | SUPER_GRC2 | 5.23 |
| **g18526** | SUPER_GRC2 | 2.68 |
| **g19121**** | SUPER_GRC2 | 49.95 |

**Table S2. Re-analysis of the pooled embryo RNA-seq data from Urban et al. (2021) (2 h–2 day embryos) identified only eight GRC-linked genes with non-zero TPM values. All eight of these genes were also detected (TPM > 0) in our own RNA-seq libraries.**
*g16029 is the only GRC-linked gene that did not show evidence of mismapping in our late-larval or adult datasets and had no highly similar paralogue on the core chromosomes. Therefore, it is the only gene considered confidently GRC-expressed in this study.
**g19121 is a GRC-linked gene in a bacterial-derived region, hence elevated expression is likely bacterial RNA contamination (see Supplementary Figure S3).

#### Supplementary Table S3: Summary BLAST results for expressed GRC-linked genes against *L ingenua* and *B. impatiens* genome

| GRC Gene ID | No. of hits | Best % identity | Best % coverage | Main Genome Hits | Comments |
|---|---|---|---|---|---|
| **g11713** | 5 | 89.7 | 100 | Bimp_SUPER_GRC, Ling_SUPER_2 | Strong cross-species match |
| **g17107** | 5 | 89.7 | 100 | Bimp_SUPER_GRC, Ling_SUPER_2 | Nearly identical to g11713 |
| **g7957** | 1 | 83 | 94 | Ling_SUPER_X | Single strong X-linked hit |
| **g8036** | 9 | 91.8 | 100 | Ling_SUPER_GRC1 | Multiple duplicated regions |
| **g13363** | 1 | 81.9 | 66 | Bimp_SUPER_GRC | Partial GRC hit |
| **g233** | 1 | 81 | 94 | Bimp_SUPER_3 | Full-length match |
| **g596** | 1 | 84.8 | 29 | Bimp_SUPER_2 | Partial alignment |

**Supplementary Table S3.** Summary of BLASTn results for expressed GRC-linked genes from *Bradysia coprophila* queried against the genomes of *Lycoriella ingenua* and *Bradysia impatiens* (from Hodson et al., 2025) . The table lists, for each GRC gene, the total number of significant BLAST hits, the highest percent identity and coverage observed, and the main genome scaffolds (SUPER contigs) containing those hits. Coverage was calculated as the proportion of the query gene length aligned in the best hit (alignment length / query length × 100). Comments summarise the strength and distribution of alignments, highlighting cases of conserved, duplicated, or partial matches across species.

The complete BLAST output files, including all alignment statistics and sequences, are provided in the accompanying Supplementary Data (under tab 11_Ling_Bimp_BLAST).
